# Late Holocene land vertebrate fauna from Cueva de los Nesofontes, Western Cuba: stratigraphy, last appearance dates, diversity and paleoecology

**DOI:** 10.1101/2020.01.17.909663

**Authors:** Johanset Orihuela, Leonel Pérez Orozco, Jorge L. Álvarez Licourt, Ricardo A. Viera Muñoz, Candido Santana Barani

## Abstract

Here we report a Late Holocene fossil-rich cave deposit from Cueva de los Nesofontes, Mayabeque Province, Cuba. The deposit’s formation and its fauna were studied through a multidisciplinary approach that included stable isotope analyses, radiocarbon chronology, stratigraphy, sedimentology, and taphonomy. Thousands of microvertebrate skeletal remains were recovered, representing a diverse land vertebrate fauna that included threatened and extinct species. The deposit is characterized by profuse *Nesophontes* remains due to raptor predation. Previously unreported last appearance dates are provided for the extinct island-shrew *Nesophontes major*, the bats *Artibeus anthonyi* and *Phyllops vetus*. Radiocarbon (^14^C AMS) age estimates between ∼1960 rcyr BP and the present were recovered. The presence of locally extinct species, including the endemic parakeet *Psittacara eups*, the flicker *Colaptes* cf. *auratus*/*fernandinae*, and the lipotyphlan *Solenodon cubanus* suggests that these species had broader distributions in the near past. Isotope analyses and faunal composition indicate the previous presence of diverse habitats, including palm grove savannas and mixed woodlands. Isotopes also provide insight into the habitat and coexistence of the extinct bat *Artibeus anthonyi* and extant *A. jamaicensis*, the diet of *Nesophontes major*, and local paleoenvironmental conditions. Oxygen isotopes reveal an excursion suggestive of drier/colder local conditions between 660 and 770 AD. Our research further expands the understanding of Cuban Quaternary extinction episodes and provides data on the distribution and paleoecology of extinct taxa. It supports the conclusion that many Cuban extinct species survived well into the pre-Columbian late Holocene and retained wide distribution ranges until human colonization.

## INTRODUCTION

Cave deposits have been, and continue to be, the richest source of extinct land vertebrate fossils in the Greater Antilles. Caves harbor different kinds of bone deposits, including accumulations due to natural death of cave inhabitants and visitors, raptor-derived pellets (e.g., mostly from owls), and dietary middens created by humans. In Cuba, these forms of bone accumulation have provided a rich vertebrate record of the island’s late Quaternary faunas, an essential source for understanding Antillean biogeography and extinctions (Morgan and Woods, 1986; Morgan, 1994; MacPhee et al., 1999).

Faunal deposits accumulated in Cuban caves were initially discovered during the mid-late 19^th^ century and the first decades of the 20^th^ century. These early efforts included discoveries by José Figueroa, Fernández de Castro, and Carlos de la Torre at several localities throughout the island between 1860 and 1911 (de la Torre, 1910; Nuñez, 1998; Goldberg et al., 2017). Later explorations were conducted by Barnum Brown (1913), Thomas Barbour, and other personnel from the Museum of Comparative Zoology (Cambridge), Carnegie Museum (Philadelphia), and the American Museum (New York City). Gerrit S. Miller (1916) and Harold E. Anthony described faunas from fossil and subfossil material found in cave deposits in eastern Cuba (Anthony, 1917, 1919), as did Peterson (1917) and Glover M. Allen in western Cuba (Allen, 1917, 1918), providing thereby the first micromammal fauna accounts from the island.

Until recently, Cuban cave fossil deposits had been rather arbitrarily considered to be of late Pleistocene age (e.g., Brown, 1913; Anthony, 1919; Allen, 1918; Koopman and Williams, 1951; Acevedo et al., 1975; Arredondo, 1970; Woloszyn and Silva, 1977; Acevedo and Arredondo, 1982; Rivero and Arredondo, 1991; Salgado et al., 1992; Balseiro, 2011). However, the few existing radiocarbon dates from non-cultural vertebrate assemblages reported from Cuba now indicate that such faunal accumulations are often much younger in age than previously expected (MacPhee et al., 1999, 2007; Jull et al., 2004; Jiménez et al., 2005; Steadman et al., 2005; Orihuela, 2010; Orihuela and Tejedor, 2012; Orihuela, 2019). So far, only three cave deposits have yielded true Pleistocene faunas: Cueva El Abrón, in Pinar del Río province (Suárez and Díaz-Franco, 2003), the tar deposits of San Felipe (Jull et al., 2004) and the thermal bath deposits of Ciego Montero (Kulp, 1952). Even though the Cuban record is one of the richest and most diverse of the Greater Antilles, it remains the least understood in terms of chronology due to the lack of reliable age estimates and discrete faunal analyses.

Such lack of chronologic resolution, which can be achieved through detailed sedimentological, stratigraphically and direct “last appearance dates” (LADs), limit our understanding of the timing of loss for most of its extinct or extirpated land vertebrate fauna. So far, of the 21 extinct land mammals, including bats, currently recognized for Cuba (Silva et al., 2007), only three, plus two birds, have direct LADs (MacPhee et al., 1999; Jull et al., 2004; Steadman et al., 2005; Orihuela, 2019). Generating additional direct and indirect LADs are crucial to constrain extinction chronologies against known past human-caused environmental changes in Cuba (Orihuela et al., forthcoming).

Here we provide a detailed, multi-proxy analysis of an exceptionally rich cave deposit from northwestern Cuba. Our interpretation of the deposit’s radiocarbon chronology, stratigraphy, and taphonomy, in addition to analyses of stable isotopes and faunal composition, contributes to the understanding of Cuban faunal diversity and biogeography by providing insight into the distribution, coexistence, diet, habitat, and timing of extinction of a wide array of taxa. The diversity and age of the deposit, plus new direct ^14^C LADs for Cuban extinct or endangered endemics, provide a unique opportunity to study a faunal assemblage that spans the critical interval between Amerindian arrival and European colonization, thus contributing to the overall understanding of Antillean land vertebrate extinction and biogeography.

## MATERIALS AND METHODS

### Geological and environmental settings

Cueva de los Nesofontes is one of a number of caves located on Loma El Palenque or Palenque Hill: lat. 23.016° N and long. −81.722° W. This hill, with a 327 m altitude, is one of the most prominent elevations of the Alturas Habana-Matanzas orographic region, in northwestern Cuba (Acevedo, 1992). Its current geopolitical position lies within the easternmost limit of Mayabeque province but was formerly included within the Province of Matanzas (Figure 1).

**Figure 1:**
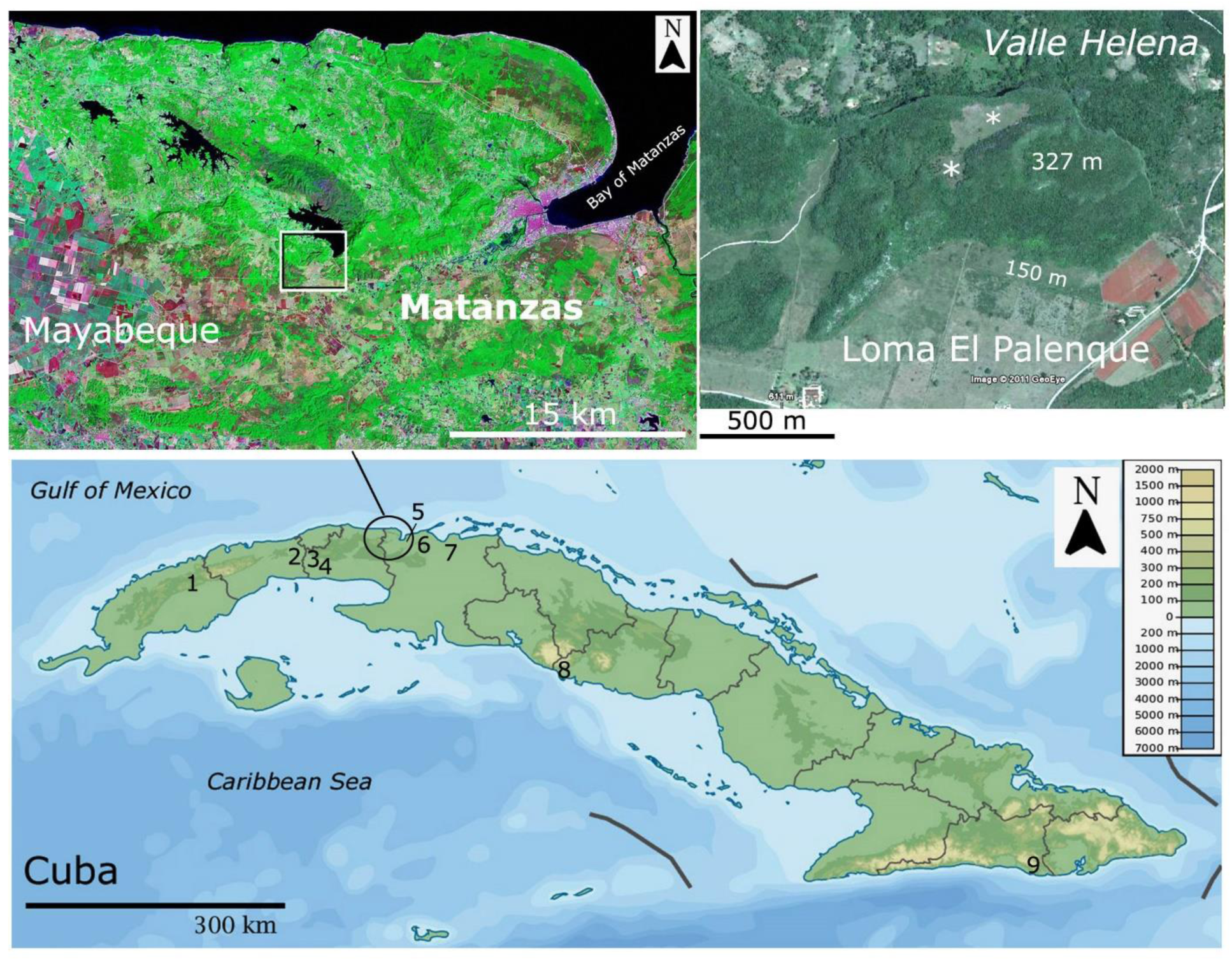
Location of Loma del Palenque and Cueva de los Nesofontes in northwestern Cuba. The asterisks (*) indicate a flat scarp at ∼260 m where red-clay soils have formed, are the main source of the allochthonous sediment inside the cave. Other important localities are indicated: **1**, Cueva El Abrón, GEDA, and Mono Fósil, Pinar del Río. **2**, Cueva de Paredones, Artemisa. **3**, Cueva del Túnel, Mayabeque. **4**, Cuevas Blancas, Mayabeque. **5**, Cueva del Gato Jíbaro, Matanzas. **6**, Cueva Calero, Matanzas. **7**, Breas de San Felipe, and Cuevas de Hato Nuevo, Matanzas. **8**, Cueva de los Masones and Jagüey, Trinidad, Las Villas. **9**, Cueva del Indio, Daiquirí, Santiago de Cuba.

Palenque is a karstic formation composed of massive (i.e., non-stratified) biodetrictic limestones of the Jaruco Fm (Formation). Previously, this hill was erroneously attributed to the Eocene (Nuñez et al., 1984; see *lapsus* in Orihuela, 2010). However, its microfauna, generally comprised of sponges, corals, mollusks, index benthic foraminifera, and echinoderms, suggest that the Jaruco Fm formed in an oxygenated, warm, tropical, neritic sublittoral-platform environment during the late Oligocene and the early Miocene, ∼28 to 20 Ma (millions of years ago) (Iturralde-Vinent, 1969a/b, 1977, 1988; Cuban Geologic Lexicon, 2014, p. 188).

Five thin sections prepared from several hand samples collected around the hill support the interpretations in the latest Cuban Geologic Lexicon (2014). The microfauna identified from those samples included large *Lepidocyclina* spp. and *Heterostegina antillea*, *Miogypsina* cf. *antillea*, and the planktonic *Globigerina* spp. *Heterostegina antillea* is an index taxon of the upper Oligocene and lower Miocene (BouDagher-Fadel, 2008). The presence of *Miogypsina* at the highest level (at 265 m above the surface of Palenque as defined by Ducloz, 1963) supports an extension for the possible formation up to the middle Miocene (JO, unp. data). As in the case of the rest of the Habana-Matanzas range, neotectonic uplift and differential erosion during the Pleistocene (< 2.6 Ma) (Iturralde-Vinent, 1988) affected the exposure of the hillside. Two of its scarp levels (the highest is indicated by asterisks in Figure 1) have been interpreted as evidence of a late Pliocene-early Pleistocene marine terrace (Iturralde-Vinent, 1969a/b, 1977), known as the Palenque Surface (Ducloz, 1963). Thus, we consider the age of the caves found within the hill to be late Pliocene or younger in age. Decomposition of exposed limestone formed the red clay ferralitic soils and loams occurring in upper escarpments (> 250 m amsl). These are known as the Matanzas red soil series (Formell and Buguelskiy, 1974), now considered as the late Quaternary Villaroja Fm (Lexicon, 2014). In terms of composition, these are the same that occur at the openings and inside of caves and fractures at Palenque.

The climate in the region is today tropical, with warm temperatures between 32 and 23 C° during the wet season (May-October), with average rainfall between ∼1300 and 1500 mm (Cuban National Atlas, 1989). During the cold - dry season (November-April) temperatures range between 18 and 26 C° (Cuban National Atlas, 1989). We registered temperatures of 6 C° inside the main gallery during the night of December 24, 2003.

Premodern vegetation was comprised of semideciduous woodlands over karst terrain and mogote forests at a higher elevation (typical mesophyll, Del Risco, 1989). Today the region is covered in secondary, but well preserved, semideciduous forest surrounded by savannas and agricultural land with lakes and rivers (Figure 1). The present vegetation on the hill includes the gumbo-limbo (*Bursera simaruba*), oaks and mahogany (*Quercus* sp. and *Swietenia* sp.), the guao (*Comocladia dodonea*), chichicate (*Urtica doica*), *Thrinax radiata* and *Coccothrinax crinita* palms and Fabaceae in the upper levels. The royal palm (*Roystonea regia*) and other agricultural plants spread through. Coffee (*Coffea arabica*) grows in the upper escarpment of the hill, and their plant remains have been observed in *Artibeus jamaicensis* roosts therein. During the colonial period, the region around the hill featured agricultural use, sugar cane, and coffee fields.

### Site-deposit description & history of research

The caves of Palenque were discovered during the late 1960s, but not fully explored or excavated until 1983–1985 by the Norbert Carteret group of the Cuban Speleological Society (Vento, 1985 in Nuñez, 1990, vol. 1: 299–304). The deposit we studied and interpret here is located inside the main gallery at Cueva de los Nesofontes, a large phreatic-vadose cave near the uppermost escarpment of Palenque (Figure1–2). The deposit is a large deposition cone situated ∼ 9 meters above the main gallery level (datum ∼ 240 m), dipping at an angle of 22–28 degrees, under a ∼ 15 meter-wide dissolution sinkhole. This sinkhole or main doline opens to other larger sinkholes with openings to the side of the hill (Figure 2). These upper caves and sinkholes are the source of the primary deposits and modern raptor roosts in which faunal remains occur or derive (Figure 2.1 and 2.3).

**Figure 2:**
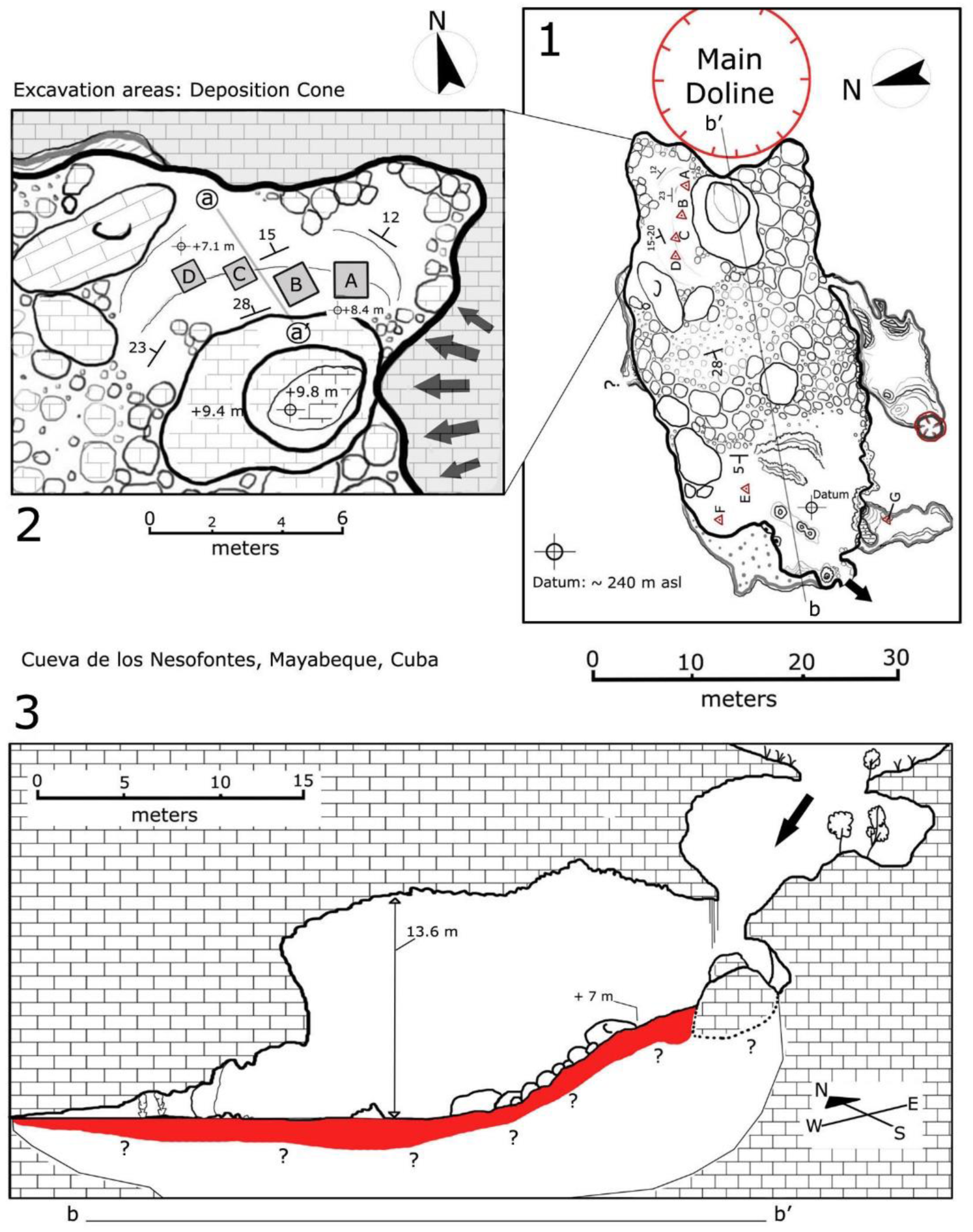
Cueva de los Nesofontes indicating geological, stratigraphic and deposition features. **1**, a gallery with main doline or sinkhole indicating areas of fauna collection: A-D pertain to the upper deposits described here. E-F are two test pits conducted on the lower level. G is the source location for the ^14^C-dated domestic dog mentioned in the text. **2**, the upper area where the main test pits described are located: A is the test pit from 1985, B-C was dug between April 1995 and December 2003. The cross-section a-a’ is the approximate sources for the stratigraphic profiles illustrated in following figures. **3**, cross-section (indicated b-b’ on **1**). Arrows indicate areas of sediment and raptor-derived pellet deposits and roost.

**Figure 3:**
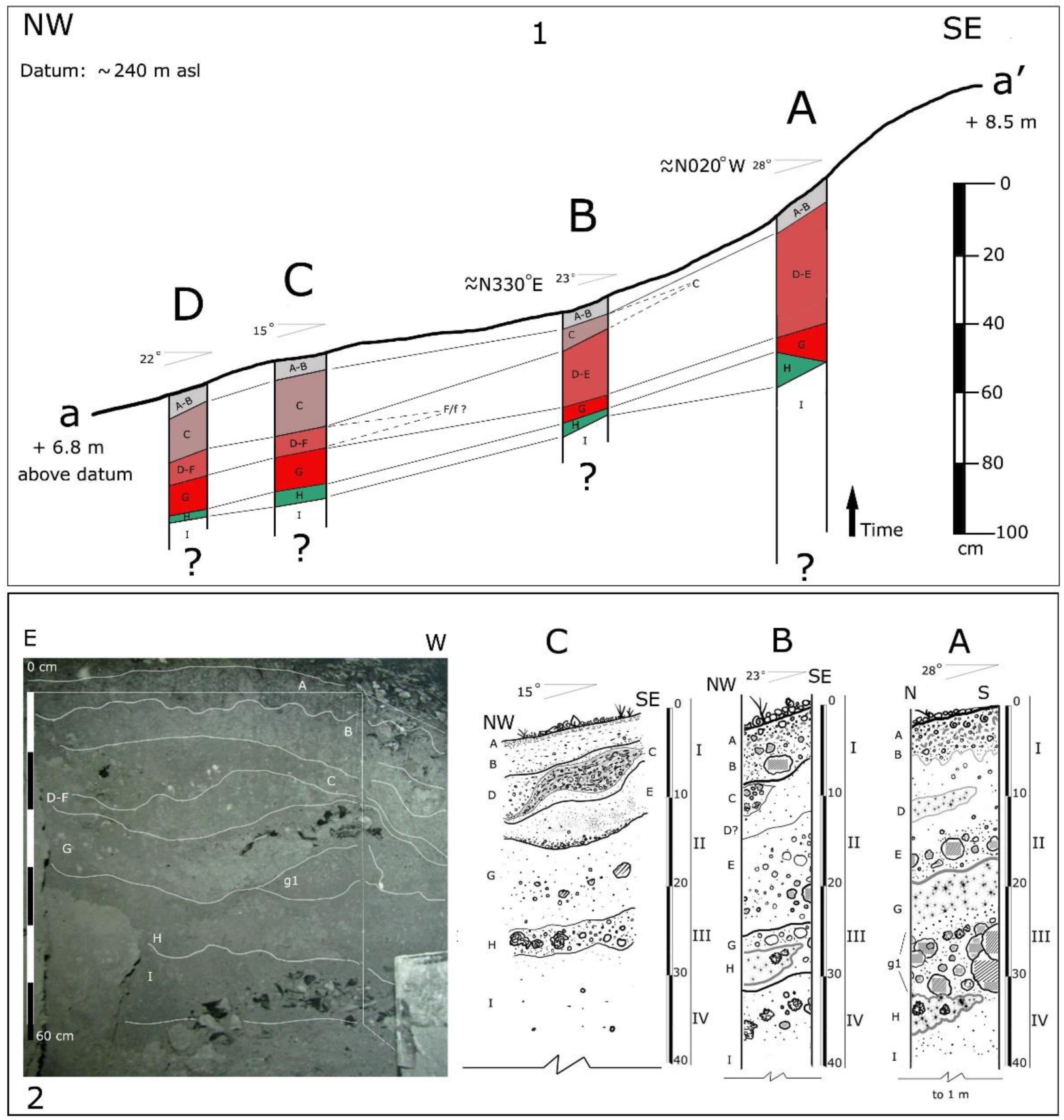
Stratigraphic profile composite, including all test pit excavations from cross-section a-a’ illustrated in Figure 2. **1**, indicates a graphic correlation of the stratigraphic units (A-I) and their lenses (e-f). The figure below shows the southern wall profile of the excavation of 1985 (A), whereas A, on the extreme lower right, shows the east wall profile. Profiles B-C pertains to the other test pits. Each Roman numeral indicates the arbitrary 10 cm intervals.

The deposit contains over 400 cubic meters of exceedingly rich fossiliferous sediment, which has been transported through the main sinkhole onto the cave’s deposition cone (Figure 2). The sediment is rill-eroded, composed of red-ferralitic soil with redoximorphic features. It is generally colored in dusky red hues and is exceptionally rich in terrestrial mollusks and *Nesophontes* remains. This abundance suggested the name of the cave as the Cave of the Island Shrew or Cueva de los Nesofontes. This cave is alternatively known as Cueva de la Caja or the Cave of the Box (e.g., Viera, 2004; Orihuela, 2019).

The main room, where the main doline and deposit are located, is littered with roof-fall boulders, smaller rocks, fallen tree branches, and leaves. The lowest level is also covered with red-colored ferralitic soil, but much less rich in biological remains. A 1.50 m test pit excavated by the Norbert Casteret group in 1985 suggests that the deposit is deeper, but not nearly as rich in fauna (Figure 3.2. and both profiles denoted A).

Although conclusive archaeological evidence has not been found in this gallery or its deposits, a ceramic fragment of unknown provenance has been recovered from the cave (Hernández de Lara et al., 2013), and a cave pictograph was recently discovered in Cueva del Campamento, situated nearly a hundred meters in the escarpment above the main sinkhole of Cueva de los Nesofontes (Orihuela and Pérez Orozco, 2015). This may relate to aboriginal or maroon occupation, as the name of the hill and the region suggests, for a Palenque is an aboriginal or maroon hideout.

### Excavation methods

Four test pits were excavated between 1985 and 2003. All excavations were done with a trowel and small metal shovel. The first and deepest test pit was excavated in 1985 (Figure 2.2 and 3) and measured 1 m length by 1m width and reached over 1 m in depth. The second had a similar measurement, but only 50 cm in depth. The last two test pits (C and D on Figure 2.1) measured 50 cm x 50 cm x 50 cm. These test pits followed 10 cm intervals with attention to the natural stratigraphy. The natural stratigraphy was identified from changes in soil coloration and faunal composition. Unconformities and erosional surfaces were detected from excavation profiles. All analyzed material was extracted *in situ* from the lateral profile into a glass vial. The data presented here originate only from test pit D.

The excavated material was dry sieved with a fine screen mesh (0.3 cm). From each sieved sample, a subsample collection was randomly placed in plastic bottles (∼ 462 cm³). This was later softly dry brushed in the lab to remove adhered matrix and soil and material separated following Silva (1974), but including juveniles and other parts of the appendicular skeleton in the tallies following the method described in Orihuela (2010). This constituted the sample collection from which species diversity was calculated.

### Stratigraphy and Sedimentology

Stratigraphic units were defined by dry color changes and changes in clast or debris size. Colors were defined using a Geologic Society of America (GSA) Geological Color Chart (2009) with a Munsell color system. The grain size was determined in the lab using USA Standard Sieves (no. 7, 2.80 mm; no. 45, 0.355 mm; no. 230, 0.0025 mm – 63 μ) placed in sequence to extract clasts from silt-clay size up to fine gravel. Percentages were calculated from bulk fraction by weight. Interval I weighed 225.7 grams; II: 30.0 g; III: 225 g; and IV: 29.8 grams. The weights were measured with an Accuris Analytical balance.

Nine levels of natural deposition (beds) were generally identified at all test pits (denoted A through I, from top to bottom). Because of the dip angle of the deposit, 2 to 3 of these beds were usually present within each of the 10 cm excavation intervals. These intervals are indicated as levels I through IV, from top to bottom. Several beds pinched out or appeared laterally as facies or lenses and are indicated with lower case letters (Figure 3).

The distinctive layers had sharp contacts with changes in coloration, which graded from the dark dusky yellow green-moderate reds of bed A and B (10 YR 4/2, 10 R 6/2 – 10 R 6/4) to the reddish oranges and moderate dusky reds (5 Y 8/4 – 10 R 6/6 – 5 R 3/4) of beds D to E. Beds were generally rill eroded, poorly sorted, with poorly rounded or subangular clasts, medium-fine sand, granules, and coarse pebbles (Table 1). Bed thickness ranged between thin and thick (5 mm to 15 cm layers). Beds A, B, G through I were near planar, wavy non-parallel, well and grade bedded, with dip angles between 22 and 28 degrees in the main slope, but less than 3 degrees at the lowest floor level of the gallery (Figure 3).

**Table 1:** Stratigraphic units, levels, and chronology from Cueva de los Nesofontes, Mayabeque, Cuba. Results and source of radiocarbon-dated (AMS ^14^C) material with lab numbers is provided.

| Interval level | Depth (cm) | Strat. Unit | Granulometry | <sup>14</sup>C Cal yrs BP | Cal yrs | Source | Number |
| --- | --- | --- | --- | --- | --- | --- | --- |
| I | 0-9 cm | A | 40.9 % med sand | 1.223 ± 0.004 pMC | 2σ: 1955-1993 AD | *Artibeus jamaicensis* humerus | ICA 15B/0116 |
| II | 10-19 cm | E | >50 % med san | 1960 ± 30 BP | 2σ: BC 40-90 AD | *Phyllops vetus* skull | ICA 18B/0845 |
| III | 20-29 cm | H | 52.4 % fine sand | 1290 ± 30 BP | 2σ: 660-770 AD | *Artibeus anthonyi* dentary | ICA 14B/1102 |
| IV | 30-45 cm | I | 49.8 % fine sand | 1418 ± 20 BP | 2σ: 605-655 AD | *Nesophontes major* dentary | Beta 392022 |
| I | 0-2 cm | n/a | Fine sand/detritus | 115.9 ± 0.6 pMC | 2σ: 1957-1993 AD | *A. jamaicensis* scapula | Beta 210380 |
| I | Surface | n/a | cave floor | 1.014 ± 0.004 pMC | 2σ: 1955-1956 AD | *Canis lupus familiaris* vertebra | ICA15B/0115 |

**Table 2:** Results of stable isotope analysis with source material from deposits at Cueva de los Nesofontes.

| Interval | Depth (cm) | Strat. Unit | δ<sup>18</sup>O<sub>apt</sub> (dental) | δ<sup>13</sup>C<sub>—</sub> | Source | Number |
| --- | --- | --- | --- | --- | --- | --- |
| I | 0 | A | -1.1 | -9.9 | *A. jamaicensis* skull | USF 15314 |
| I | 0 | A | **n/a** | -20.1 col. | *A. jamaicensis* humerus | ICA 15B/0116 |
| I-II | ~14 | C | -1 | -8.1 | *A. jamaicensis* humeri | USF 15316 |
| II | ~18 | E | -0.9 | -10 | *A. jamaicensis* skull | USF 15313 |
| III-IV | 29-31 | H | -0.7 | -11.0 | *A. anthonyi* dentary | USF 15315 |
| III-IV | 29-32 | H | **n/a** | -21.1 col. | *A. anthonyi* dentary | ICA 14B/1102 |
| IV | >32 | I | -1 | -20.7 col. | *N. major* dentary | Beta 392022 |
| I | 0-2 cm | n/a | **n/a** | -20.7 col. | *A. jamaicensis* scapula | Beta 210380 |
| I | Surface | n/a | **n/a** | -10.9 col. | *C. lupus familiaris* vertebra | ICA15B/0115 |

The beds were separated by sharp contacts or boundaries (i.e., disconformity/erosional surfaces), especially between beds C, D, E, and F. Layers A, B, and G–I were generally conformant or paracomformant (i.e., of undiscernible unconformities). Bed C constituted a large first-order ash bed with fragments of charcoal, wood detritus, coarse clasts, abundant fossils and gastropod shells (ash made up > 30 % composition). This layer contained exotic species such as murids and the domestic European sparrow (*Passer domesticus*). The beds H – I formed the largest paracomformity with unidentifiable layers below the ∼ 50 cm depth (Level IV) (Figure 3– 4).

**Figure 4:**
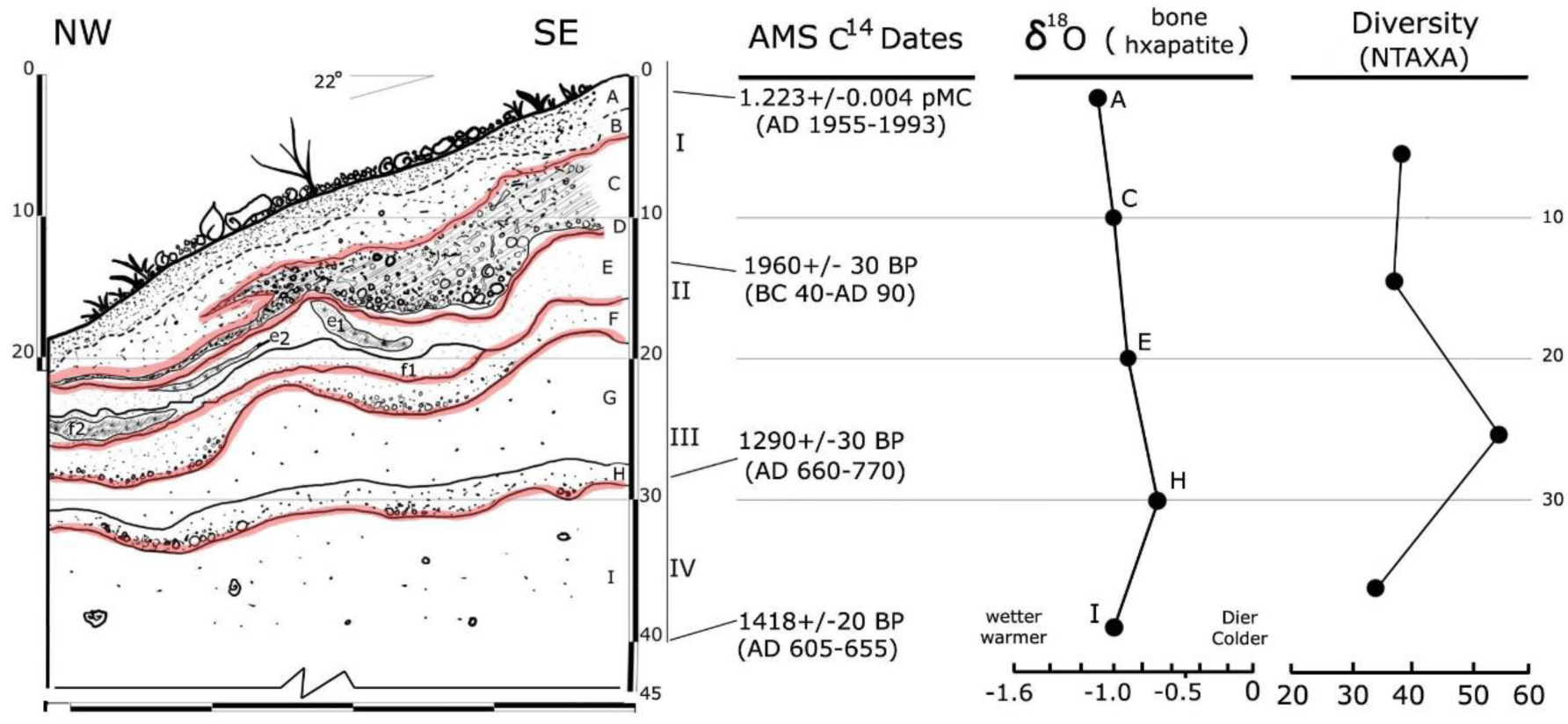
Lateral profile of test pit D with source radiocarbon dates, oxygen stable isotopes and NTAXA diversity by intervals. The fauna and multi-proxy analyses described in the text are from this excavation. The red lines indicate the major disconformities/erosional surfaces.

Most beds were correlated between test pits (Figure 3). Others, such as bed E, F, and G included small lenses (e1, e2, f1, f2, and g1), that graded laterally or pinched out up-slope. Bed C also pinched out towards the higher parts of the deposition zone, where H also seemed to disappear, at least laterally (Figure 3.1, 3.2).

### Multifaceted Analytical approaches

For elemental analysis, high-resolution imaging, and characterization of cave soils and loams we used a JEOL JSM 5900LV scanning electron microscope (SEM) with energy dispersive spectroscopy EDS-UTW with detectors of 3.0 nm resolution at the Florida Center for Analytical Electron Microscopy (FCAEM) facility at Florida International University (Miami, FL). Soil or fossil fragments selected for analysis were placed in separate stages, and each sample analyzed three times. The averages are reported in weight percentage (wt %) of those measurements. These analyses allowed for the identification of clay particles, other clasts content, and the overall elemental composition of the red clay soils. These analyses were conducted without coating, directly on dry samples kept in sterile glass vials collected *in situ*. For microscope and thin-section analysis, a Leica DM EP petrographic microscope was used. The samples were prepared at Florida International University.

Radiocarbon dating ^14^C AMS (accelerator mass spectrometry) and several of the isotope analyses (for nitrogen and carbon) were conducted by Beta Analytic Inc. (Miami, FL), and International Chemical Analysis Inc. (ICA, Ft. Lauderdale, FL), following each lab’s standard procedure and who reported no complications (D. Wood, R. Hatfield, and B. Díaz, pers. Comm. 2014-2018). The dates and most isotope values were determined from bone collagen. These are reported using the standard notation of radiocarbon years before the present (rcyrs BP). Carbon younger in age than the modern reference standards is reported as “Percent Modern Carbon” (pMC), which indicate a date after thermonuclear testing, and date after the 1950s (Hua and Barbettii, 2004).

The conventional ^14^C AMS dates were calibrated to calendar age-intercept solar years (Cal. yrs.) to one and two sigma ranges (±1σ −2σ) using Oxcal v4.3, on IntCal13 carbon curve for the Northern Hemisphere (Reimer et al., 2013). See also Ramsey (2017) at https://c14.arch.ox.ac.uk/oxcalhelp/hlp_contents.html. Only values that differed less than 140 years were considered contemporaneous (Semken et al., 2010), although the rule of thumb may extend up to ±200 years due to multiple intercepts and conversion curve topography on dates during the last 2000 years (Geyh and Schleicher, 1990 in MacPhee et al., 1999). Late Quaternary epochs and time intervals discussed follow Morgan and Woods (1986), Soto-Centeno et al. (2015) and limits established by the IUGS (International Union of Geological Sciences).

Additional isotope analyses were conducted at the Stable Isotope Ratio Mass Spectrometry Facility at the University of South Florida (USF, Tampa, FL). These analyses were conducted to explore paleoenvironment and diet that could be interpreted from isotope signals (Bocherens et al., 1996; Ben-David and Flaherty, 2012). Such additional data could help elucidate aspects of competition and habitat selectivity between some of the species analyzed.

Carbon (C), oxygen (O) and nitrogen (N) isotope values were determined from bone apatite and collagen and their rations reported in delta (δ) standard notation: ^13^C/^12^C = δ¹³C_apt. for carbon acquired from apatite and δ¹³C_col. when acquired from bone collagen. The same applies to nitrogen: ^14^N/^15^N= δ^15^ N_apt. (apatite) and δ^15^ N_col. (bone collagen). The carbon from apatite is reported in parts per mil (‰) compared to the Vienna Pee Dee Belemnite (VPDB) and nitrogen from atmospheric nitrogen (AIR) (Ambrose and Norr, 1993; Bocherens et al., 1996). Oxygen values, ^18^O/^17^O =δ ^18^O, were acquired from tooth apatite of *Artibeus jamaicensis* remains, and are reported also as a ratio of VDPB parts per mil (‰). These values likely originate from available drinking water or water in the fruits consumed by the *Artibeus* bats, and thus provides a regional paleoclimatic proxy (Bocherens et al., 1996; Ben-David and Flaherty, 2012). The C: N ratio used to indicate diagenesis or alteration in the collagen sample was always below 3.4, suggesting insignificant or no diagenesis on the analyzed remains (DeNiro, 1985; Bocherens et al., 1996; Ben-David and Flaherty, 2012).

### Taphonomic and fauna methodologies

The weathering levels, based on a numerical value representative of bone erosion, flaking or fracturing due to atmospheric exposure follow Behrensmeyer (1978), Shipman (1981) and Andrews (1990). Criteria for bioturbation index follows Tylor and Goldring (1993). Estimation of taxonomic abundance, diversity and their indices follow Lyman (2008).

Anatomical terminology for birds follows Howard (1929), Olsen (1979) and for mammals Silva et al (2007). Systematic taxonomy of Cuban rodents follows Silva et al (2007). For *Nesophontes* we follow Rzebik-Kowalska and Woloszyn (2012) and our work in preparation in considering three valid species in Cuba. The validity of *Nesophontes micrus* and *N. major* are furthermore supported by proteomics, despite the inherent limitations of this analysis (Buckley et al. submitted). For extant Cuban birds, we followed Garrido and Kirkconnell (2000), González (2012), and for extinct birds, Orihuela (2019) and others cited in the text.

Fauna and faunal variations discussed here only pertain to test pit D. We infer that Pit D does not differ from the others, which were slightly less diverse, but similarly rich in *Nesophontes* spp (Author’s unp. Data). Tables 3 and 4 provide a synthesis of the fauna present in the Pit D assemblage. Moreover, Table 4 provides a stratigraphic distribution of taxa within each of the levels and beds of Pit D. The fauna we will discuss ahead pertain to only species which are noteworthy or represent extralimital records.

**Table 3:**
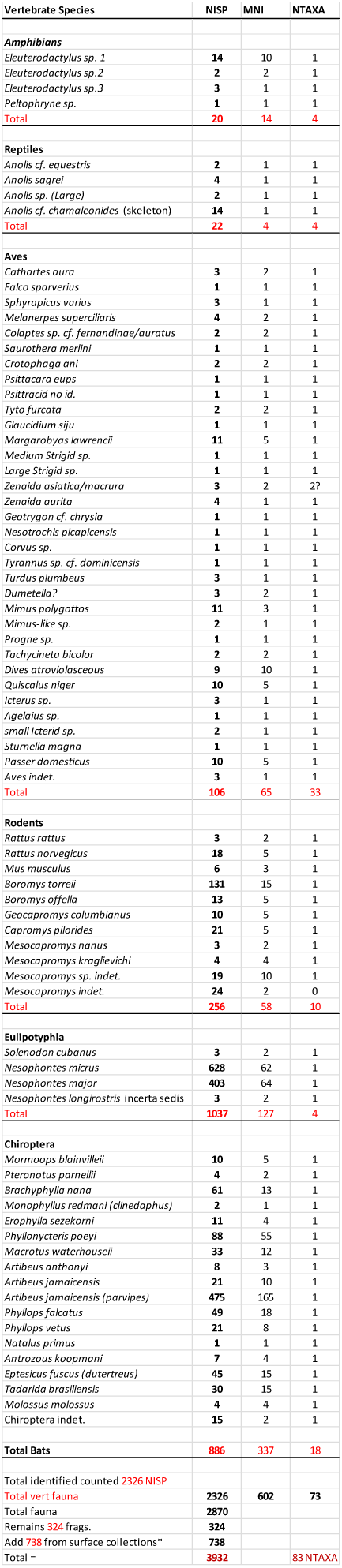
Cueva de los Nesofontes fauna list, providing a number of individual specimens (NISP), the minimum number of individuals (MNI), and the number of total taxa (NTAXA) counts.

**Table 4:**
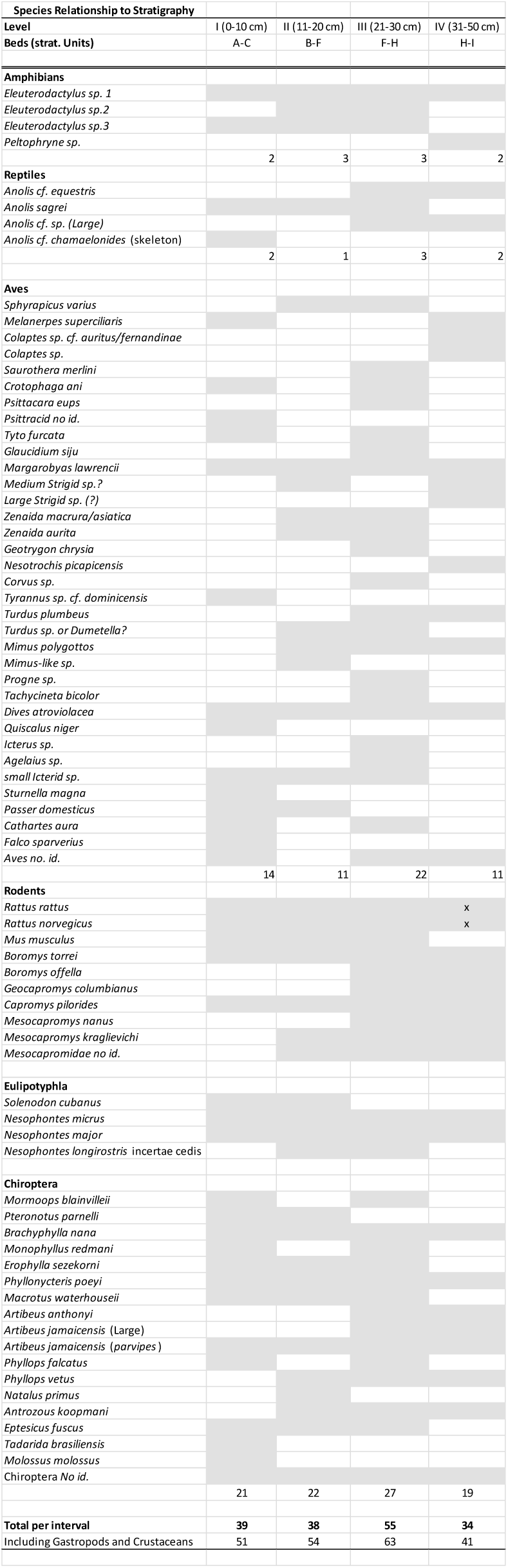
Stratigraphic distribution of taxa throughout each interval and their individual count. Grey-filled boxes indicate the presence and empty absence. A total by interval is provided.

Specimens were compared and identified with neontological and fossil collections at the American Museum of Natural History (AMNH), in New York City (USA), the Museum of Natural History of the University of Florida (UF-FLMNH) in Gainesville, Florida (USA), the Institute of Ecology and Systematics (IES), La Habana (Cuba), and zoological collection of Gabinete de Arqueológia, Office of the Conservator of the city of La Habana, Cuba. All the remains analyzed were extracted with permission of the Central Registry of National Cultural Goods (certification nos. 20141965; LHA–23, fol. 0162773). All the remains from these and other excavations are deposited in the collection of the Museo Nacional de Historia Natural (MNHNCu), in La Habana, Cuba. Part of the collection has been cataloged (Donation 13.18: MNHNCu–72–05.01 and 76–156–215), but the rest remains uncatalogued (E. Aranda, persn. Comm. 2016, 2018).

Measurements were taken with a digital caliper and are reported in millimeters (mm). All statistical analyses were conducted with the software PAST v3 and STATISTICA software (1995, v5). Two-way ANOVAs and Tukey’s Test for unequal sample sizes were used to compare linear measurements between species. Principal component analysis (PCA) was performed to further explore differences between *Nesophontes* taxa, and the first two extracted principal components were used to generate a plot. Probabilities were compared to a significance level of alpha < 0.05, and of <0.01 for the PCA. These data were plotted using STATISTICA (1995).

## RESULTS

### Radiocarbon Chronology and sedimentation rates

Four radiocarbon dates (^14^C AMS) were acquired from the four stratigraphic intervals of test pit D (Table 1; Figure 4). For the upper level (I), a fresh *Artibeus jamaicensis* adult humerus was selected from bed A. For Level II, a skull of the extinct bat *Phyllops vetus* from bed E. From lowermost (near interface) level III, a dentary of the extinct bat *Artibeus anthonyi* from bed H, and for level IV, a dentary of the extinct shrew *Nesophontes major* from bed I. These last three radiocarbon dates represent the first direct LADs reported for these Cuban species.

The uppermost bed (A) yielded a modern carbon age between 1955 and 1993 AD, and thus a very modern age for this level. The date for bed E, between BC (BCE) 40 and 90 AD (CE) revealed an inversion event in the stratigraphy or reworking of older remains since the lower levels yielded younger dates between AD 605–655 and AD 660–770 (Figure 3; Table 1).

An additional date was acquired for a domestic dog (*Canis lupus familiaris*) skeleton found mineralized in the floor of a small room at the entrance of the doline gallery (Figure 2.1, collection site G; Table 1). Originally, this specimen was considered Amerindian in age and was thus selected for testing. However, the age it yielded indicated its deposition within the modern period AD 1957–1993 and is likely contemporaneous with bed A of the cone deposit above. A similar surface radiocarbon date from this cave, albeit a different deposit, is provided in Orihuela (2010). All these superficial tests help support that the uppermost levels of the cave’s deposit are generally modern (i.e., post-Columbian). But the presence of extinct taxa such as *Nesophontes* there too, suggests likely partial reworking.

All dates suggest ample hiatuses of several hundred years between beds/intervals (Figure 4). These had slow sedimentation rates that varied between 1.15 mm/yr at the upper level (beds A–C), and slightly faster rates > 1.30 mm/yr for the middle levels (bed C–E), and 1.28 mm/yr for the lower III-IV, beds H and I.

### Stable isotopes

Stable isotopes of carbon (δ¹³C) and oxygen were measured from apatite (δ¹³C_apt.) and bone collagen (δ¹³C_col.) of four adult specimens of the fruit bat *A. jamaicensis*, plus one adult specimen of the extinct bat *A. anthonyi* and a newborn *Canis lupus familiaris* (the same which were ^14^C dated; Table 1). Moreover, oxygen and carbon isotopic values were acquired from four *A. jamaicensis* dental apatite samples from each interval (Table 2).

An additional analysis of nitrogen (δ^15^N_col.) and carbon (δ¹³C_col.) isotopes were obtained from the bone collagen of the ^14^C dated *N. major* (Table 2). This specimen yielded a value of −20.7 ‰ δ¹³C_col. and of 7.9 ‰ δ^15^N_col. These data help approximate the diet of these vertebrates and provide insight into the paleoenvironments and taphonomy, as are interpreted in the Discussion section.

### Taxon identification and fauna sample

A total of 3932 specimens were collected from the assemblage (test Pit D), of which 2326 (59.2 %) were identifiable vertebrate specimens (NISP) and 324 were unidentifiable fragments. The NISP increased to 2870 if invertebrates were included (Table 3). Another 738 specimens were collected from two other surface deposits within the cave near the deposit (Figure 2). The total, including invertebrates, represented 83 taxa (NTAXA).

Of the total NTAXA (*n*=83), 73 taxa represented vertebrates, yielding a count of 602 minimum number of identified individuals (MNI) (Table 3). This fauna was mostly composed of birds (33 species) and mammals (∼32 species), 39.8 % and 38.6 % of the total NTAXA respectively. Of the birds, the woodpeckers (at least 3 taxa or 9 %), the strigids (at least 3), pigeons (at least 3) and passerines (7 or 21%), were the most abundant.

Within the mammals, the bats and lipotyphlans were the most abundant, but the rodents and bats were the most diverse (Table 3). NTAXA diversity increases to 77 if other species records from the surface collections and other excavated deposits within the cave are added. These include the bats *Desmodus rotundus*, *Chilonatalus macer* and *Lasiurus insularis* (Orihuela, 2010).

The gastropod fauna was diverse with at least 9 species preliminarily recorded. Further identification of their remains will likely result in an increase in overall NTAXA count. The gastropods, amphibians, and reptiles will not be discussed in detail here. These groups of organisms have been poorly studied in Cuban Quaternary deposits, and thus our knowledge of them in the recent past is very limited. In the case of the amphibians and reptiles, this has been largely dictated by a lack of modern comparative osteological material in the Cuban zoological collections (Aranda, 2019). However, those that we could identify (Table 3) will be briefly commented on in the Discussion, and altogether add to the knowledge of the island’s past herpetofauna.

### Species Accounts: noteworthy or extralimital record fauna

Aves

Accipitriformes

Cathartidae Lafresnaye, 1839

*Cathartes aura* (Linnaeus, 1758)

**Material**: one left femur (MNHNCu uncataloged, field no. 582a) and a complete skull (MNHNCu uncataloged, field no. 582b) without mandible from bed A (level I), and one incomplete premaxilla (MNHNCu uncataloged, field no. 193) from bed G (level III) (Figure 5.1). A complete skeleton with evidence of anthropogenic combustion was found at the lower part of the main doline gallery, but not collected.

**Figure 5:**
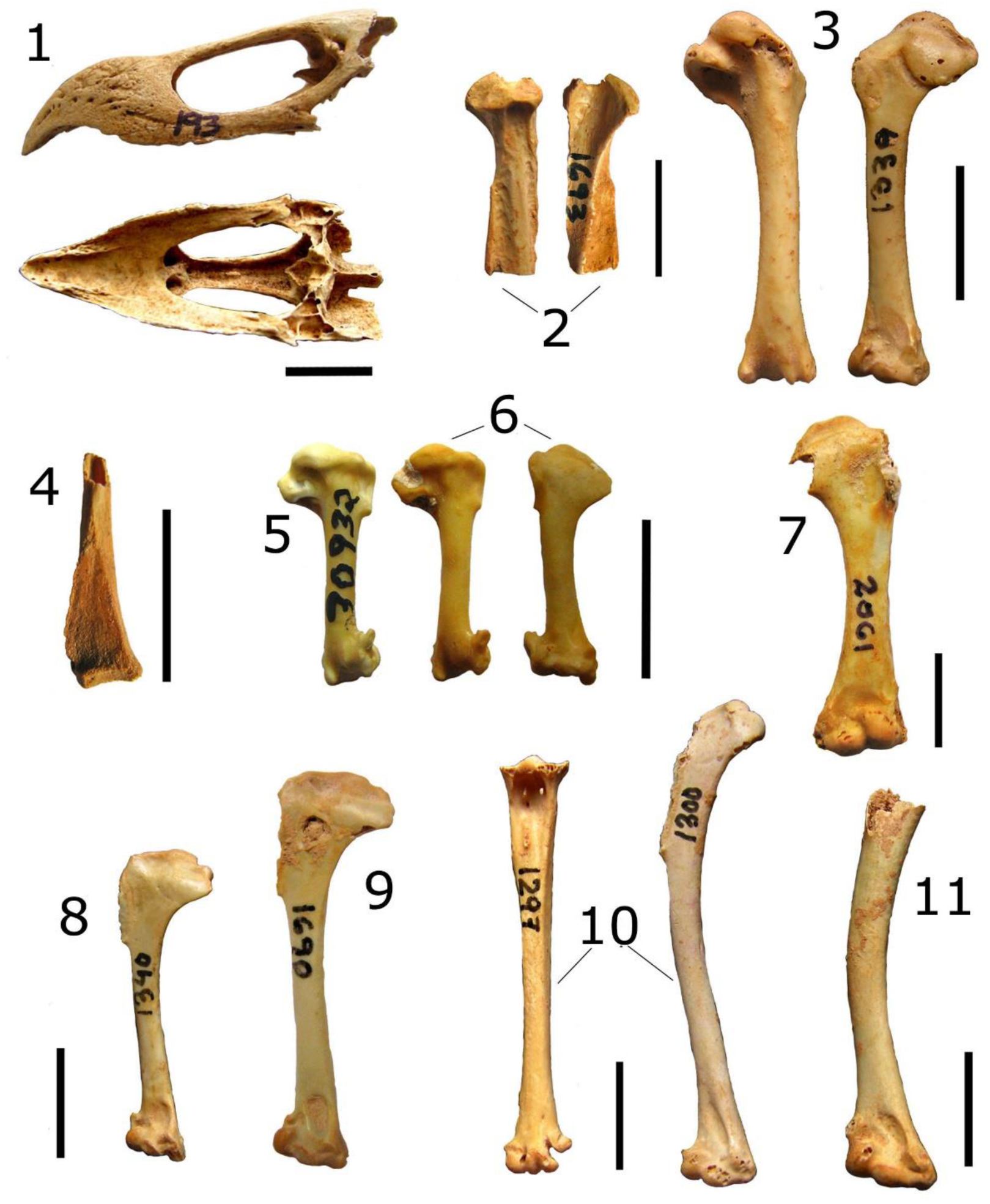
Aves. Fossil and subfossil bird remains from test pit D. 1, *Cathartes aura* maxilla in lateral and dorsal view. 2, proximal tibiotarsus of *Colaptes* cf. *fernandinae*. 3, the humerus of *Psittacara eups*. 4, distal coracoid of *Progne* cf. *cryptoleuca* or *subis*. 5, humerus of *Tachycineta bicolor* (FLMN 17685). 6, humerus of *Tachycineta* cf. *bicolor*. 7, humerus of *Geotrygon* cf. *chrysia*. 8, humerus of *Sphyrapicus varius*. 9, humerus of *Melanerpes superciliaris*. 10, tarsometatarsus (left) and humerus of *Margarobyas lawrencii*. 11, humerus of *Saurothera merlini*. Each scale bar represents 10 mm.

**Description**: with the fossil fragment provided in parenthesis, the specimens measured as follows: maximum skull length 91.9 mm, maximum upper maxilla length 51.2 mm (45.2 mm), maximum nasal opening width 18.1 mm (17.1 mm), and maximum maxillary width 14.3 mm (14.5 mm). The femur measured in maximum length (GTL) 69.4 mm, proximal maximum width (GPW) 18.9 mm, distal maximum width (GDW) 17.6 mm, and a maximum width of the diaphysis (shaft-GSW) 18.1 mm. The fossil premaxilla is not mineralized but showed slight evidence of corrosion and weathering.

**Taxonomic remarks**: Suárez (2001) mentioned the existence of two undescribed extinct vultures from Cuba. One of them is apparently referable to *Cathartes* but is not *C. aura* (Orihuela, 2019). However, the specimen reported here seems indistinguishable quantitatively or qualitatively from *C. aura* (Figure 5.1). Our specimen from layer G lacks a direct date, but it was found between the dated contexts ranged between 1690±30 and 1290±30 rcyr BP and is thus preliminarily considered late Holocene/pre-Columbian in age. This, therefore, constitutes the first pre-Columbian record of the species in Cuba.

Piciformes

Picidae Leach, 1820

*Colaptes* sp. cf. *fernandinae* (Vigors, 1827) or *auratus* (Linnaeus, 1758).

**Material**: a single, distal tibiotarsus fragment from layer G (level III) (MNHNCu uncatalogued; field number 1693) (Figure 5.2).

**Description**: This is a weathered specimen with evidence of digestion. It measures in greatest distal width (GDW) 5.01 mm and in greatest shaft width (GSW) 2.2 mm.

**Taxonomic remarks**: This specimen is slightly larger than *Melanerpes superciliaris* (uncatalogued from this deposit), *M. radiolatus* (UF 27075), GDW 4.90–4.91 mm and GSW 1.93–1.95 mm, and *Xiphidiopicus percusus* (UF 36476: GDW 4.06 mm and GSW 1.6 mm). About similar size or slightly smaller than *Colaptes auratus* (UF 45035: GDW 5.67 mm and GSW 1.96 mm), which suggests a medium-sized woodpecker (∼ 33–35 cm; Short, 1965). In Cuba, the only two woodpeckers that fall within this size category are the endemic Fernandina’s flicker *Colaptes fernandinae* (∼ 34 cm) and the flicker *C. auratus* (∼ 33 cm) (Garrido and Kirkconnell, 2000). Our tibiotarsus specimen (no. 1693) resembles *Colaptes* more than *Melanerpes* in having marked and narrower intermuscular line and low (unflattering) fibular crest. The outer cnemial crest is more arched or circular in our specimen, as in *Colaptes* and not more open as in *Melanerpes*. However, we did not compare it directly to *C. fernandinae*, and thus its identification remains tentative. An additional proximal tibiotarsus (no. 1794) from layer I (level IV) is similarly attributed to this taxon (O. Jiménez pers. Comm. 2015, 2018).

Psittaciformes

Psittacidae Rafinesque, 1815

*Psittacara eups* (Wagler, 1832) sensu Remsen et al. (2013).

**Material:** A complete right humerus (field number 1339) from layer G (level III) (Figure 5.3).

**Description:** Well preserved specimen, measuring in total length (TL) 28.2 mm, GDW 5.8 mm, greatest proximal width (GPW) 9.26 mm and GSW 2.69 mm.

**Taxonomic remarks:** This specimen compares in size with *Psittacara* parakeets such as *Psittacara nana* from Jamaica (UF 25929): TL 29.8 mm, GDW 6.01 mm, DPW 10.1 mm and GSW 2.55 mm. Morphologically is most similar to this genus in having a shallow bicipital furrow, scarcely grooved bicipital furrow and deltoid crest, round head, poorly developed external tuberosity proximally. Distally, shallow brachial depression and etepicondylar prominence. It was qualitatively comparable to the endemic Cuban parakeet *P. eups* (Garrido and Kirkconnell, 2000). This specimen was associated with the species aforementioned, and are likely of the same age. This constitutes the first pre-Columbian record for the species.

Passeriformes

Hirundinidae Rafinesque, 1815

*Progne* sp. cf. *cryptoleuca* (Gmelin, 1789) or *subis* (Linnaeus, 1758)

**Material:** Incomplete, distal left coracoid, stained brown red (field number 1624), from layer H (level III).

**Description:** This specimen may represent a juvenile because of its porosity and rounded sternal facet (Figure 5.4). Measurements: GDW 4.39 mm and GSW 1.75 mm.

**Taxonomic remarks:** This coracoid represent a swallow larger than any other of the species present in Cuba. In morphology, it is similar to *P. subis* but slightly smaller. The purple martin (*P. subis*) and the Cuban martin (*P. cryptoleuca*) are common in Cuba. The first is a common transient between August and March, whereas the second is a common resident nearly year-round (Garrido and Kirkconnell, 2000, p. 168). Neither species has been previously reported from the paleontological or Amerindian record of Cuba.

Hirundinidae Rafinesque, 1815

*Tachycineta* cf. *bicolor* (Vieillot, 1808)

**Material:** A complete left humerus (MNHNCu, uncatalogued) from layer G (level III).

**Description:** The specimen is slightly mineralized, small and delicate. It measures in GTL 15.3 mm, GDW 5.5 mm, GSW 1.6 mm, and GPW 6.6 mm. (Figure 5.6).

**Figure 6:**
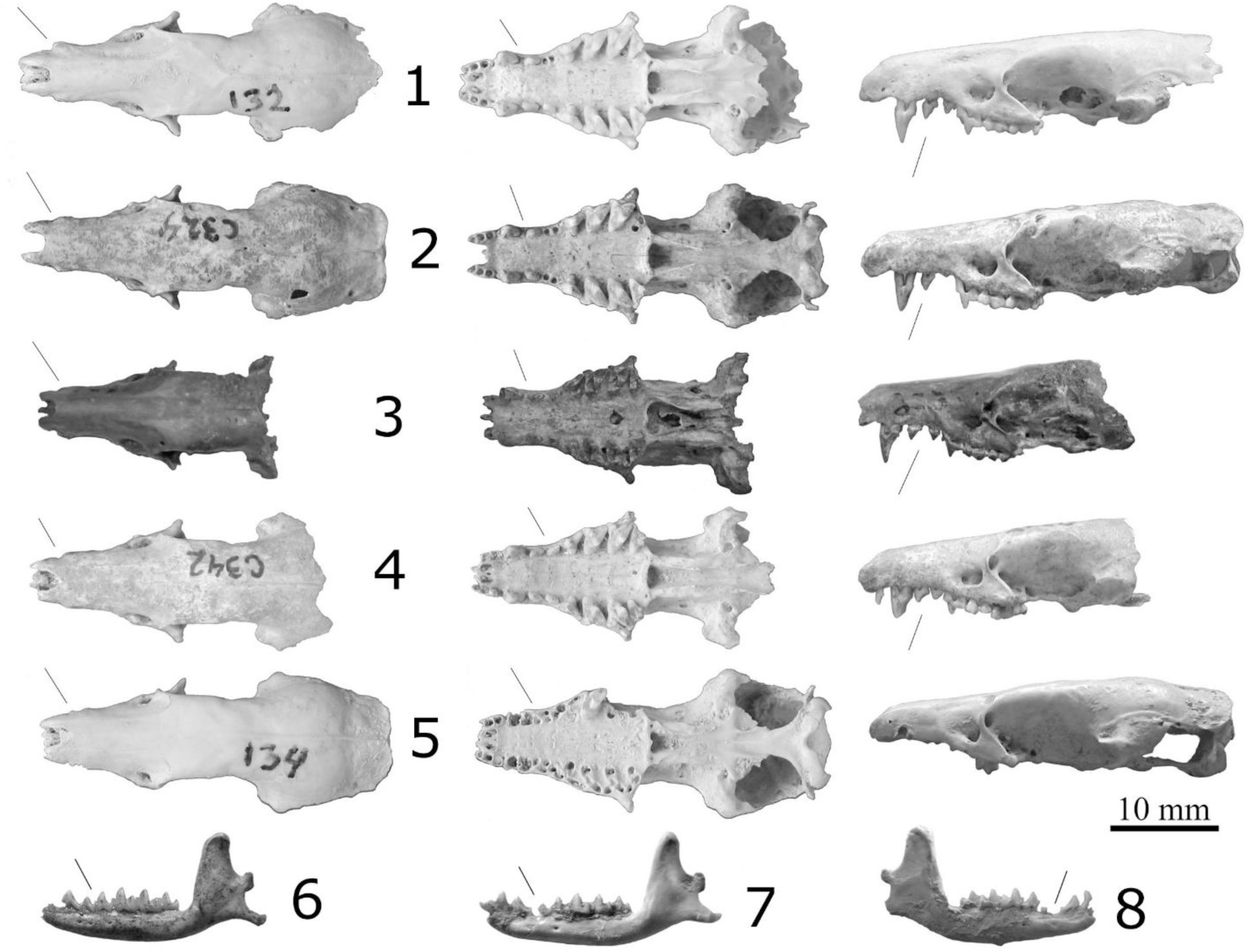
Comparison of tentatively identified *Nesophontes* cf. *longirostris* specimens (1–3 skulls, 3 is the holotype AMNH 17626, 7-8 tentative mandibles) and *Nesophontes major* (4–6). Small lines indicate discrete characters discussed in the text. Note the more elongated rostrum and wider gap between upper premolars in *N. longirostris* and the crowding in *N. major*.

**Taxonomic remarks:** This specimen is remarkably similar to the tree swallow *T. bicolor*, a common transient in Cuba (Garrido and Kirkconnell, 2000, p. 169). Our specimen agrees well in size and morphology to a male from Indian River, Florida, USA (UF 17685/30932): GTL 15.3–15.4 mm, GDW 4.91–5.22 mm, GSW 1.62–1.64 mm, and GPW 6.44 mm (Figure 5.5). The ectepicondylar prominence is prominent and grooved at the tip, with a slight lateral extension (rome, shorter and attached in *Hirundo rustica* and hook-like in *Progne subis*). The internal condyle entepicondyle is less pronounced than the external condyle, but more than the intercondylar furrow, which is slightly flattened (not in *H. rustica* or very pronounced in *P. subis*). The bicipital furrow and deltoid crest are poorly developed off the main shaft. The capital groove is deeply excavated, unlike *Hirundo*, which has a double furrow (deep single furrow in *Progne*). Thus, we refer it tentatively here to *T. bicolor*. A direct comparison to the Bahamian tree swallow *T. cyaneoviridis* was not conducted. However, this taxon is a slightly larger rare winter transient in Cuba (op. cit.). This represents the first paleontological and prehistoric record for Cuba.

Mammalia

Rodentia

Capromyidae Smith, 1842

*Mesocapromys* Varona, 1970

**Material:** This genus is represented by over 50 specimens, most of which are long bones, representing at least 2 species and 20 individuals. The two species are represented by *Mesocapromys nanus* and *Mesocapromys kraglievichi*. This genus was present at all levels and in most beds, but more profusely in level III and IV (Table 4).

**Description:** Most remains showed taphonomic evidence of predation and digestion. Others were mineralized or adhered to a carbonate matrix. Most were juveniles with open or incomplete epiphysis and alveoli.

**Taxonomic remarks:** Although Silva et al. (2007) and M. Condis (unp. Data) provided size groups for elements of the appendicular skeleton, attributing any of these long bones to a specific species is problematic due to lack of complete skeletons as comparative material. Often, identification and assignment are satisfactory when complete adult hemimandibles are present in the assemblage, for which there are diagnostic *M. nanus* and *M. kraglievichi*. At present, the only diagnostic trait distinguishing them is the lateral extension of the condyle’s ascending ramus process beyond the plane orientation of the angular process in *M. nanus* when the dentary is in occlusal view (i.e., viewed from above; Silva et al., 2007 p. 176). In *M. kraglievichi*, the ascending ramus follows the same plane as the angular process below. Most of the undetermined material assigned to *Mesocapromys* spp. indet. Table 3 represents juveniles, just as those of the extinct *Geocapromys columbianus* and the extant *Capromys pilorides*, which were well-represented in the assemblage (Table 3–4).

Lipotyphla

Solenodontidae Gill, 1872

*Solenodon cubanus* Peters, 1861

**Material:** A left proximal ulna fragment from layer E (level II). A complete edentulous right mandible (uncatalogued) and complete left scapula (MNHNCu, field no. 2029) from a surface collection near the deposition cone and under the main sinkhole. This last specimen yielded a direct ^14^C age of 650±15 BP (UCIAMS 218808; Orihuela et al., forthcoming).

**Description:** The surface specimens likely belong to the same individual, and appeared fresh (weathering level 0), with slight discoloration. The ulna was slightly mineralized and showed evidence of cracking (weathering level 1) and represents another individual from the sinkhole deposit above.

**Taxonomic remarks:** These specimens are indistinguishable from *Solenodon cubanus*. The radius was associated with the bat *Phyllops vetus* that yielded an age of 1960 rcyr BP, thus indicating a pre-Columbian, late Holocene age for that specimen, whereas those from the surface may be several hundred years old, as is supported by the ^14^C age estimate of the left scapula (no. 2029).

Nesophontidae Anthony, 1916

*Nesophontes* sp. cf. *longirostris* (sensu Anthony, 1919)

**Material:** Three specimens may represent this taxon: a near-complete skull, lacking the occipital and petrosals (MNHNCu field no. 132), and two possible hemimandibles (MNHNCu, field no. 121 and 1428). The first skull and mandible are from layer E (level II), and the last (no. 1428) was from layer H (lower level III).

**Description:** Large species of *Nesophontes*, like *N. major* (Figure 6.4–6.6), but with a tubular and more elongated rostrum, wider diastemata between upper and lower canine and first two premolars. Skull 132 and dentary 121 were slightly mineralized, and dentary 1428 partially mineralized. Measurements provided in Table 5 and plot graphs in Figure 7.

**Figure 7.**
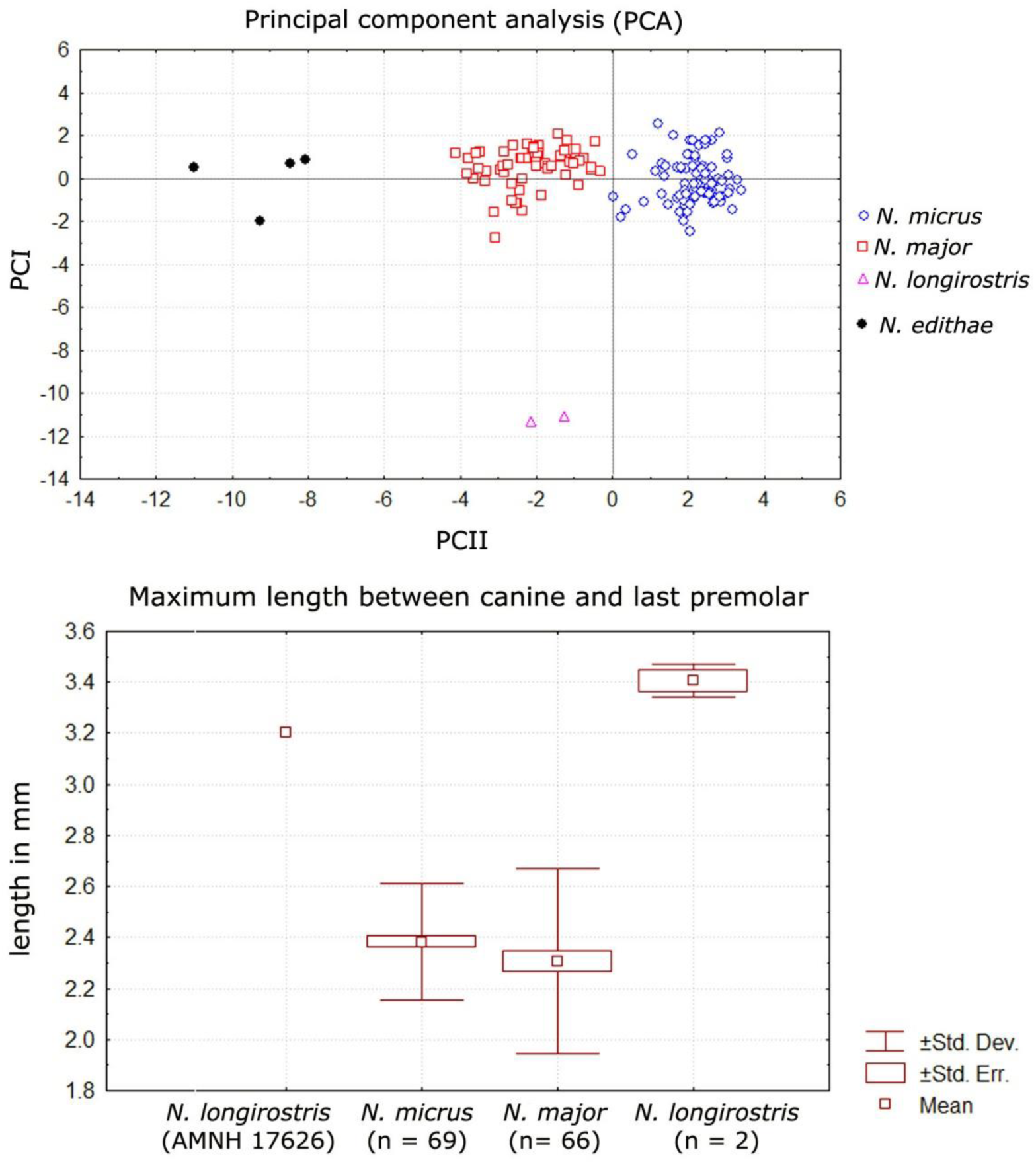
Plot graph of principal component analysis (PCA), above, shows the loading of root 1 and 2 for several *Nesophontes* species. Note the distinctive separation of *N. longirostris* from *N. major*, *N. micrus* and *N. edithae*, used as an outgroup, (p < 0.10). Plot graph from 2-Way ANOVA with Tukey’s post hoc test (p < 0.05), indicating the quantitative range of variation in the maximum length between posterior canine to anterior third maxillary premolar in *N. micrus*, *N. major*, *N. longirostris* from Pit D, and *N. longirostris* holotype.

**Table 5:** Nesophontes craniomandibular measurements, including the specimens tentatively identified here as *N. longirostris*. All specimens are from Cueva de los Nesofontes, except two *N. longirostris* (one is the AMNH holotype, and the other a specimen from Cueva del Gato Jíbaro referenced in the text). * = p < 0.050.

| Species | Measurement (mm) | N | mean±SD | min-max |
| --- | --- | --- | --- | --- |
| *N. micrus* | Max. Dental length | 56 | 12.89±0.3 | 12.2-13.56 |
|  | C-M3 length | 67 | 10.55±0.3 | 10.01-11.33 |
|  | Max. Palatal length | 56 | 13.09±0.4 | 12.2-13.99 |
|  | Max. Canine breath | 68 | 3.61±0.2 | 3.23-4.66 |
|  | Max. Canine-Pm3 length | 70 | 2.35±0.2 | 1.7-2.91 |
|  | Max. dentary length | 243 | 18.03±0.6 | 16.0-19.3 |
|  | Max. dentary dental length | 257 | 11.48±0.3 | 10.18-12.56 |
|  | Max. c-pm3 length | 6 | 3.07±1.7 | 2.66-3.49 |
| *N. major* | Max. Dental length | 43 | 14.53±0.6 | 12.63-15.46 |
|  | C-M3 length | 56 | 11.75±0.5 | 10.02-12.65 |
|  | Max. Palatal length | 42 | 14.7±0.5 | 13.56-15.53 |
|  | Max. Canine breath | 62 | 4.86±0.04 | 4.23-5.44 |
|  | Max. Canine-Pm3 length | 65 | 2.30±0.04 | 1.59-2.98 |
|  | Max. dentary length | 147 | 20.77±0.7 | 18.09-22.6 |
|  | Max. dentary dental length | 160 | 13.0±0.5 | 11.44-14.4 |
|  | Max. c-pm3 length | 9 | 2.92±2.1 | 2.31-3.45 |
| *N. longirostris* | Max. Dental length | 4 | 14.87±1.2 | 13.58-15.83 |
|  | C-M3 length | 4 | 12.35±0.09 | 11.80-12.4 |
|  | Max. Palatal length | 4 | 15.43±0.6 | 14.0-15.9 |
|  | Max. Canine breath | 4 | 4.85±0.4 | 4.4-5.26 |
|  | Max. Canine-Pm3 length* | 4 | 3.33±0.1 | 3.20-3.45 |
|  | Max. dentary length | 2 | 22.34±0.3 | 22.08-22.60 |
|  | Max. dentary dental length | 2 | 13.72±0.2 | 13.54-13.90 |
|  | Max. c-pm3 length | 2 | 4.0±0.4 | 3.58-4.42 |

The skull of *N. longirostris* is most like that of *Nesophontes major* (Figure 6) but differs in being slightly larger, with a slenderer and more elongated rostrum, more parallel postorbital, with a wide diastema between the upper canine and the first two maxillary premolars (Pm1-Pm3). The is also a wider separation between the last incisor and the canine. In *N. major*, the rostrum is broader, more U-shaped, and wider at the level of the canines. The angle of inclination of the nasal is more pronounced in *N. longirostris* than *N. major*.

*N.* longirostris shows an incipient tapering at the level of the first and second maxillary premolars not present in *N. major* (including juvenile individuals). The orientation and size of the premolars in *N. micrus* are nearly parallel to the axis of the toothrow and of nearly equal size. In *N. major*, the premolars are always crowded, oriented obliquely from the toothrow, and the first premolar is always larger than the second. In *N. longirostris*, the orientation of the premolars is slightly oblique, despite their wide separation. In *N. longirostris* the Paracone is reduced in the third upper molar (M3) but is smaller and slimmer than M1 and M2. M1 is slightly smaller than M2 and very subtriangular in shape. In *N. major* the M3 is more robust and wider (more quadrate), with a slightly higher Paracone, and the M1 is stubbier than the M2, with a less pronounced Metastyle (Figure 6).

In this sense, *N. longirostris* seems more akin to *N. major* than to *N. micrus*. Quantitatively, the two species are also most similar in most cranial linear measurements. *N. longirostris* is slightly larger in skull, palatal and dental length, likely as a function of the wider spacing between the premolars. In maximum length taken from the posterior canine to the anterior premolar defined by Anthony (1919), they are significantly larger (p = 0.000736) than *N.* major and *N. micrus* (Figure 7). In this measurement, they are even larger than the holotype of *N. edithae*.

The dentary of *Nesophontes major* (both supposed males and females) are significantly (p < 0.050) larger than *micrus* in several linear dimensions: total length of the dentary 20.8 (18.09-22.6), *N. micrus* 18.0 (16.0-19.3); maximum height of coronoid process 10.0 (8.64-11.33), *N. micrus* 7.71 (6.6-8.46); and maximum height of the mandibular ramus under m1-m2 3.09 (2.36-3.74), *N. micrus* 2.27 (6.6-8.46). In general, the dentary and lower dentition of *N. major* is more robust and marked than *N. micrus*. The dentary of *N. major* has a thicker ramus, with a more pronounced curve at the masseteric/digastric region (thinner, and much less curved in *N. micrus*; the muscle scar is less pronounced). The shape of the coronoid process is wider, broader, with more pronounced masseteric fossa on the lateral face, and deeper temporalis/pterygoid fossae on the medial face (subtriangular, thinner, less marked or shallow, and more restricted in *N. micrus*). The canine of *N. major* is an ungrooved premolaliform, with a small cingulum and more triangular cusp and smaller base (wider base and wider triangular-wider shear surface outline in *N. micrus*). In the molars, the angle between the paraconid and metaconid, as seen on lateral aspect, is more closed, with a wider commissure (more open and lower in *N. micrus*, with a reduction in cingulum development). The scar of the mandibular symphysis in *N. major* is more pronounced and longer than in *N. micrus*. In this sense, the supposed mandible of *N. longirostris* is nearly identical to *N. major*, but with the diastemata present between pm1 and pm2. Based on this qualitative and quantitative, *N. longirostris* is tentatively revalidated here but will be further discussed elsewhere.

**Taxonomic remarks:** H. E. Anthony described this species based on an incomplete skull (AMNH 17626; Figure 7.3) from a cave deposit in Daiquirí, southeastern Cuba. He distinguished it from *N. micrus* by its longer and more slender rostrum, plus a “distinct diastemata between the canine and the first premolars” (Anthony, 1919, p. 634). Anthony also predicted that such diastema would be found in the dentary. This diastema resulted in a larger measurement of 3.2 mm taken between the posterior border of the maxillary canine and the anterior border of the premolar, in comparison to other specimens he studied (op. cit.). Since Morgan (1977) and subsequent revisors considered *N. longirostris* invalid and a synonym of *N. micrus* (Condis et al., 2005; Silva et al., 2007; Rzebik-Kowalska and Woloszyn, 2012). Despite these evaluations and considering the intra and interspecific variation of the genus (JO pers. Obs.; Buckley et al., in pub.), the characters displayed by these specimens seem to suggest otherwise.

Our specimens, both skulls, and dentaries, have the supposed diagnostic diastemata, elongated rostrum and measurements that exceed the observed variation in both *N. micrus* and *N. major* studied from multiple locations in Cuba (*n* > 720 hemimandibles and *n* >150 skulls; plus over 1030 specimens from this assemblage alone) and Anthony’s Daiquirí series at the AMNH. Moreover, adding the discovery of another complete skull specimen (MNHNCu, field no. 324; Figure 6.2) with similar morphology and measurements from Cueva del Gato Jíbaro, ∼18 km east from the assemblage described here. This last specimen is associated with the archaeological kitchen midden dated to 860±30 BP (Orihuela et al., forthcoming).

Chiroptera

Phyllostomidae Gray, 1825

*Artibeus anthonyi* Woloszyn and Silva, 1977

**Material:** eight specimens (MNHNCu, uncataloged), representing at least three individuals in the assemblage belong to this species. These were a rostrum, three hemimandibles (no. 11, 12, and 1663), and four humeri encountered within layer H (lower level III) and layer I (level IV) (Figure 8.1).

**Figure 8:**
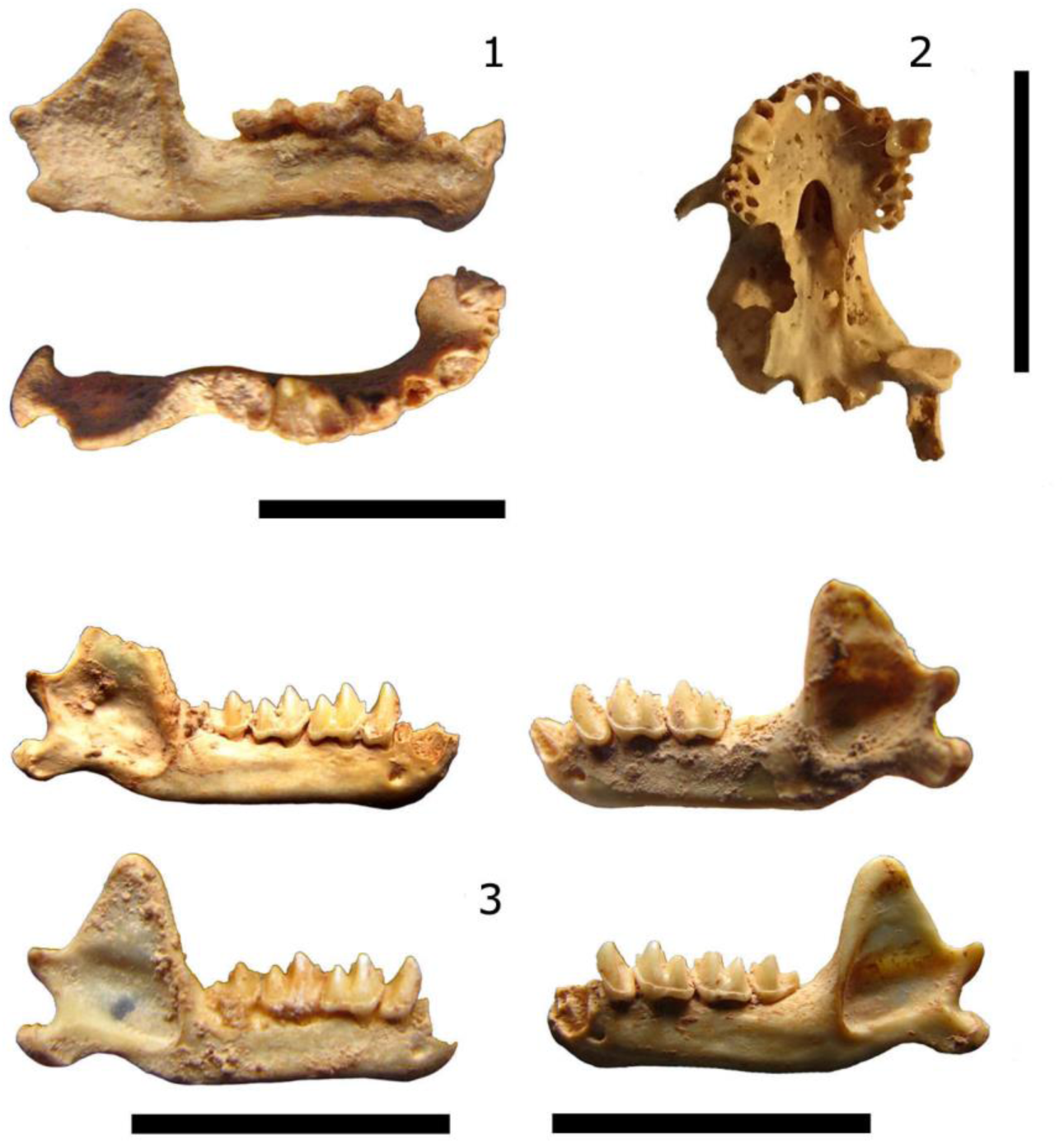
Bats. 1, left hemimandible of *Artibeus anthonyi* (no, 1663, lower Level III). 2, the incomplete skull of *Phyllops vetus*, on the ventral view (no. 37, Level II). Both specimens radiocarbon-dated (^14^C). 3 hemimandibles of *Antrozous koopmani* (no. 20, 75, 1429, 1430 from Levels II and IV, also ^14^C dated). The scale bar = 10 mm.

**Description:** These specimens were mineralized, with a few including calcareous encrustations. One of them, a slightly mineralized and robust right hemimandible (no. 1663) found at the bottom of layer H (lowermost level III) yielded a direct radiocarbon date of 1290±30 rcyrs BP (Figure 8.1), providing the first direct LAD for this taxon in Cuba.

**Taxonomic remarks:** The humeri measured between 36.0 and 37.7 mm, and the mandibles had a total length greater than 18.4 mm and less than 22.0 mm. These specimens were identified from *Artibeus jamaicensis*, and the Cuban subspecies *parvipes*, based on size and criteria published by Anthony (1919), Woloszyn and Silva (1977), Silva (1979), Balseiro et al. (2009) and Orihuela (2010). *Artibeus anthonyi* has been reported from another deposit in Cueva de los Nesofontes (Orihuela, 2010). The species seems to have been widespread in the archipelago. So far, *A. anthonyi* has been documented from 11 localities (Borroto-Páez and Mancina, 2017).

Including this record and another from a paleontological layer at Cueva del Gato Jíbaro adds to 13 localities. This last specimen yielded a middle Holocene ^14^C direct date estimate (Orihuela et al., forthcoming).

*Artibeus jamaicensis* Leach, 1821

**Material:** The Jamaican fruit bat was represented by 173 skulls, 254 mandibles, and 45 humeri. Radii and other parts of the appendicular skeleton were not fully counted, but more than 22 specimens, including scapulae and femora, represented this species. NISP of 495 and an MNI of at least 165 individuals (Table 3).

**Description:** After *Nesophontes micrus* and *N. major*, this taxon was the third most common taxon of the assemblage. Remains of this species displayed multiple taphonomic marks of deposition, mineralization, decomposition, predation, and digestion (see Figure 10.6).

**Figure 10:**
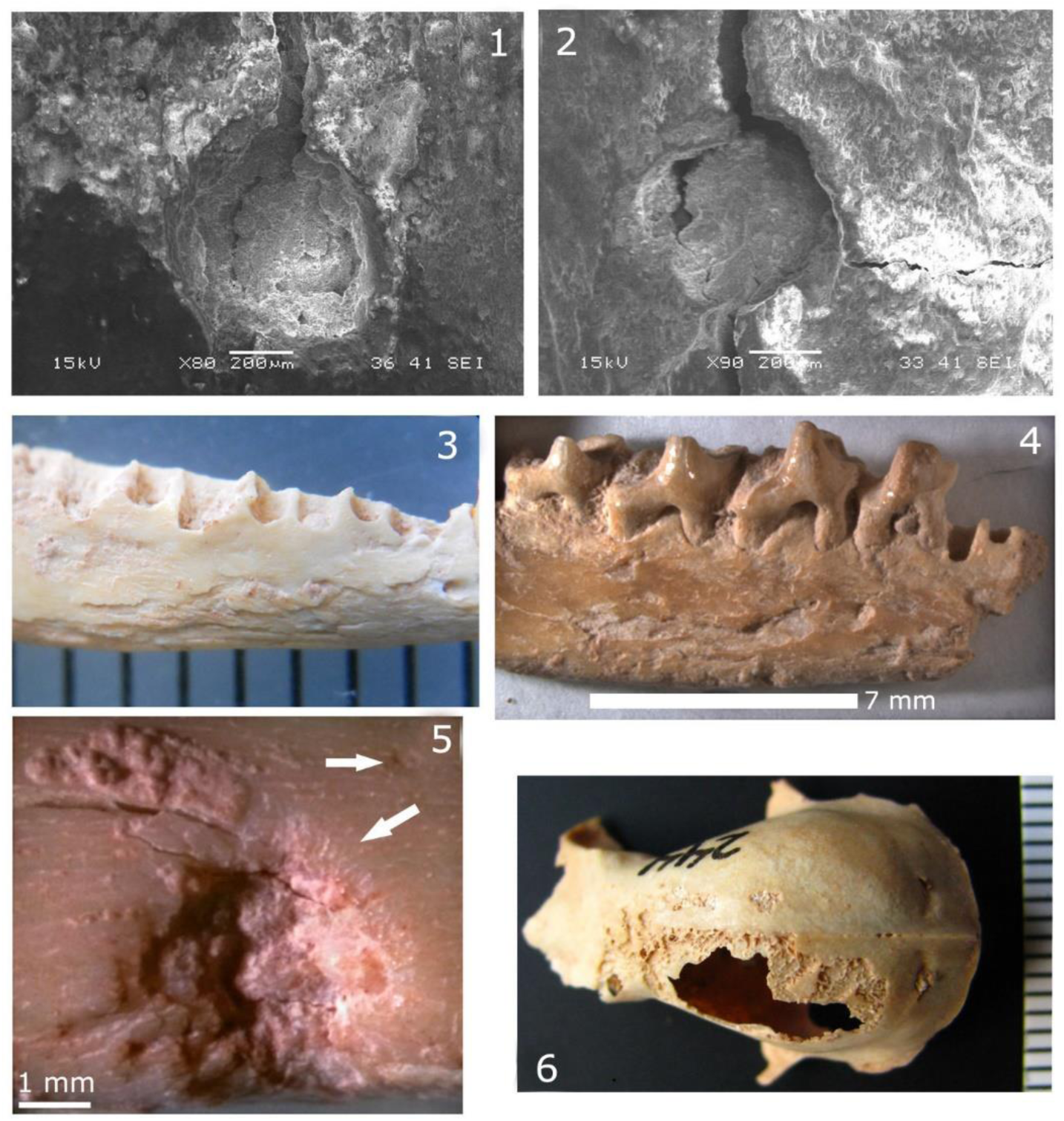
Taphonomic evidence. 1-2, images are SEM microphotographs of *Nesophontes* sp. tooth marks on small capromyid long bones. Note the well-rounded edges, microfractures radiating from the cortex. 3-4, show stage 3 weathering on an *N. major* left hemimandible (left) and stage IV on another (right). 5, shows microscopic striae and notching associated with insect scavenging on the bone. The main depression is likely a tooth mark made on the bone while still fresh (note the gradual peeling features). 6, is an *A. jamaicensis* adult skull with evidence of raptor predation and digestion (corrosion).

**Taxonomic remarks:** The majority of these specimens are indistinguishable morphologically and metrically from the Cuban endemic subspecies *A. jamaicensis parvipes*. However, eight crania, eight hemimandibles and four humeri (NISP of 21), indicated in Table 3 as *A. jamaicensis*, were larger than the maxima of the fossil and neontological range provided by Silva (1974, 1979) and Balseiro et al (2009). These specimens slightly exceeded the upper range of *A. jamaicensis parvipes* in palatal length (> 13.5 mm), anteorbital width (> 8.5 mm), and postorbital breath (> 7.2 mm) (Silva, 1979). In this last measurement, it also exceeded values reported for *A. anthonyi* (> 7.4 mm; Woloszyn and Silva, 1977; Balseiro et al., 2009) and *Artibeus lituratus* (> 6.7 mm in Woloszyn and Silva, 1977). This variation may be a form of temporal or chronoclinal variation but will be further explored elsewhere. Since these specimens are qualitatively inseparable from *A. jamaicensis*, they are included within this taxon. These specimens occurred exclusively in layers H and I (levels III and IV) where they were directly associated with *A. jamaicensis*, *A. anthonyi*, and *Phyllops vetus*.

*Phyllops vetus* Anthony, 1917

**Material:** Taxon represented by eight fragmentary skulls, including rostra, nine dentaries, and three humeri, representing at least eight individuals (MNHNCu, uncataloged).

**Description:** Most remains were fragile and slightly mineralized. A skull (no. 37) found in layer E (level II; Figure 8.2) yielded a radiocarbon age of 1960±30 rcyrs BP, constituting the first direct LAD for this species.

**Remarks:** This taxon appeared in association with the Cuban fig-eating bat *P. falcatus* only in layer G (level III), which yielded radiocarbon ages between ∼1960 and 1290 rcyrs BP; Table 4. *P. vetus* occurred in all levels except level I (layers A–D, in Figure 4). These age estimates are further supported by radiocarbon dates now available for this level (Orihuela et al., forthcoming).

Vespertilionidae Gray, 1821

*Antrozous koopmani* Orr and Silva, 1960

**Material:** This taxon was represented by a partial skull (MNHNCu uncataloged), a fragmentary braincase (MNHNCu uncataloged) and five dentaries (MNHNCu uncataloged, field no. 19, 20, 75, 1429, 1430), occurring in all layers between level II and IV (Figure 8.3). Three of these have provided direct radiocarbon dates from beds F, G, and I, that agree with the overall Late Holocene age estimates for these intervals (Orihuela et al., forthcoming).

**Description:** The specimens were well-preserved, often showing evidence of predation and digestion. They did not deviate quantitatively or qualitatively from other reported specimens (Orr and Silva, 1960; Silva, 1976; 1979; García and Mancina, 2011). Viera (2004) reported other specimens from surface collections in the same cave.

**Taxonomic remarks:** The Cuban pallid bat is in need of a detailed revision. Although it is often considered a subspecies of the continental species *Antrozous pallidus* from western North America (Simmons, 2005), we consider that the differences in morphology and size warrant its retention as a distinct endemic species until further analyses are conducted (following Silva 1976; Silva and Vela, 2009; García and Mancina, 2011). This species was undetected in Cuba until the mid-20th century. The first, and only complete specimens preserved were two females collected by Charles T. Ramsden in 1920–21, near Bayate, Guantanamo, eastern Cuba, but misidentified as “*Macrotus*” (Silva, 1976). *A. koopmani* has been found in several “fresh” owl pellets across the island, which suggest a former wide range in the island, but has not been confirmed captured or observed live since 1956 (Orr and Silva, 1960; Silva, 1979; Borroto-Páez and Mancina, 2017), although a questionable report exists (see comm. in Mancina, 2012). Moreover, MacPhee and colleagues have shown that pellet material that is apparently “fresh” can be several hundreds of years old (1999). This species is extremely rare in collections, currently extremely endangered or already extinct.

### Other organisms

Pollen, plants seeds, phytoliths, and starch grains were detected at all intervals of the deposit but remain unstudied (Figure 9).

**Figure 9:**
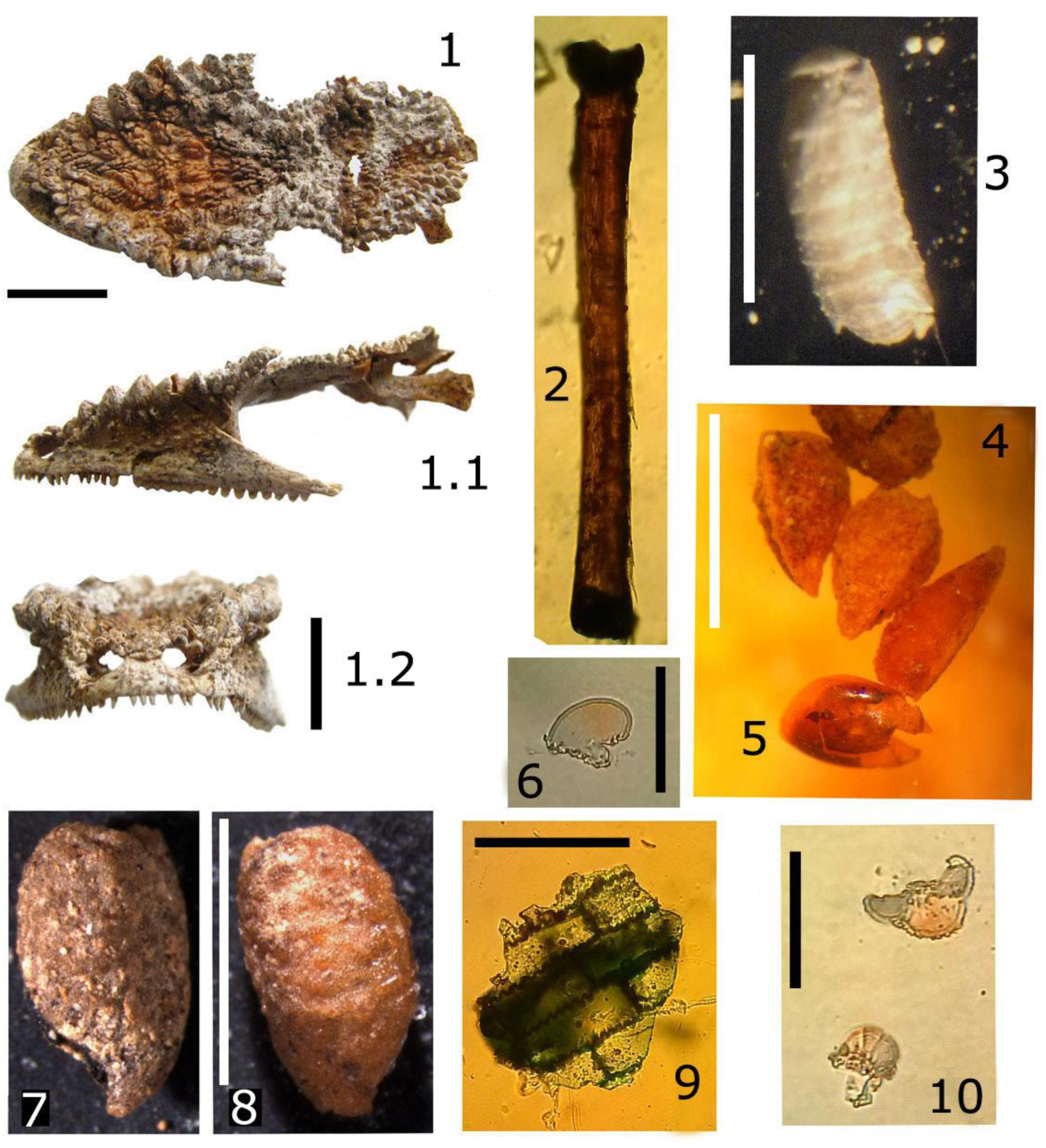
Plant, invertebrate and reptile remains. 1-1.2, *Anolis* cf. *chamaleolis* (no. 606), scale bar 10 mm (Level I). 2, unidentified insect extremity ∼ 1 mm (Level IV). 3, photid fly pupa, scale 10 mm (Level IV). 4-5, plant seeds, scale 5 mm. 6 fungus spore (?), scale ∼ 50μm (All from level II and III). 7-8, seeds, scale 5 mm. 9, leaf fragment, scale ∼ 50μm (Level II). 10 plant spores, likely a conifer, *Pinus*? (Level IV), scale ∼ 50μm.

Gastropods and crab remains were very common throughout the deposit. At least nine species of land snails and a land crab, *Gecarcinus ruricola*, were present and abundant in the assemblage. The land snails included the following preliminary taxa: *Alcadia* sp. cf. *hispida*, *Farcimen* cf. *procer*, *Chondropoma* cf. *vespertinum*, *Oleacina subulata*, *Opisthosiphon* sp., *Nescoptis* sp., *Liguus fasciatus,* and *Zachrysia auricoma*. The last two and *Chondropoma* sp., being the most abundant. Unidentified plant fragments such as leaves, bark, microcharcoal, and seeds were also present (Figure 9.4-9.9).

Insect chitin was present in the matrix of the upper levels (I and II). Within the lowest levels, microscopic fragments of insect exoskeletons and fly pupae were rare but well preserved when present (Figure 9). One of the pupae specimens was identified as a phorid fly pupa (Figure 9.3). Remains of larvae were observed directly on the bones of several specimens at the level I and III.

Amphibians were represented by at least two genera, *Eleutherodactylus,* and *Peltophryne* spp, but otherwise difficult to assign to species. The Cuban tree frog *Osteopilus septentrionalis* is likely also present. The reptiles were identified as lizards of the *Anolis* group: the smaller *Anolis sagrei*, the larger *Anolis equestris*, a similar large *Anolis* sp., and *A.* cf. *chamaeolonides* (fide Nicholson et al., 2012; Rodríguez-Schettino et al., 2013), this last on Figure 9.1.

### Taphonomic observations

Mineralization, coloration, and evidence of predation and digestion were the most common taphonomic evidence (Figure 10). Weathering was another important factor acting on the preservation of the specimens. Evidence of predation in form of scratches, claw or beak marks, indentations, fractured braincases, and digestion corrosion, were much more frequent in the upper levels (I and II), whereas most mineralization and maximum weathering levels (> level 2) were more evident in lower levels. Weathering levels or stages varied generally between 0 and 2, only rarely did specimens show stages higher than or equal to 3 (Figure 10.3, 10.4). Scavenging evidence in the form of gnawing and tooth marks by rodents and *Nesophontes* island-shrews (Figure 10.1, 10.2) has been documented in detail from this assemblage (Orihuela et al., 2016).

Decomposition-related insect activity such as boreholes, etchings, and fungal activity was less common (Figure 9.5), but likely related to the exposure of the pellets before and during the formation of the deposit. In several cases, the soft clay of the deposit invaded the empty braincase cavities of several *Nesophontes* specimens, creating natural endocasts (Orihuela, 2014).

Skulls and mandibles were the most common of all skeletal elements, with 476 and 1359 specimens respectively; they contributed 14.2 % and 59.1 % to the osseous remains in the assemblage (Pit D). Thus cranial elements, especially mandibles, dominated the assemblage at 79.8 %. Humeri (133 specimens) represented 4%, and other elements of the appendicular skeleton (398 specimens), likely constituted a total of 17.2%. It is important to note, however, that many radii and femora were fragmented and unidentifiable to species level, and thus, not counted.

### Pathologic observations

Evidence of pathologies was present in less than 1 percent of the assemblage. These were evident in the bats *Artibeus jamaicensis*, capromyid rodents, and *Nesophontes*, in the form of bone lesions, healed fractures, general bone deformations, and dental-alveolar lesions. Three specimens of *Nesophontes major* were of special note: A left adult dentary showed a markedly open premolar root with indications of an alveolar infection. Two other hemimandibles showed, as supported by radiography (not illustrated here), healed fractures or deformed coronoid processes. Mineralization, insect activity, and digestion often caused corrosion on the bones that could be mistaken for fungal or pathologic conditions (Figure 10.5).

## DISCUSSION

### Source of the fossils: Sedimentology and interpretation of deposit formation

The vertebrate fossils that compose this assemblage presumably mostly originated from raptor-derived primary pellet deposits located above the main sinkhole that was slowly in-washed (transported) into the cone of deposition under the sinkhole. Based on the faunal composition of the upper layer and surface samples collected around the deposit, we can infer that other organisms were included in the assemblage also from natural death, such as the crustaceans, gastropods, reptiles and several birds and bats. Among the samples collected from isolated non-pellet deposits included *Canis*, *Tyto* and *Cathartes* aforementioned, plus an articulated skull and mandible of *N. micrus* found on a nearby wall. All these suggest other sources for fauna in the deposit.

With the organic remains came sediments from the upper scarp levels of Palenque Hill. Based on the SEM-EDS data, these soils were positively correlated (R²=0.8353; y=0.4526x + 1.9158) in Si, Fe and Al weight percent composition with ferralitic clay soils of the Mayabeque-Matanzas lowlands (Formell and Buguelskiy, 1974), and with the ferralitic-ferromagnesic red soils of the upper scarp of Palenque Hill (asterisks in Figure 1). The changes in coloration are redoximorphic features, indicating depletion of oxidizing/reducing Fe-Mn conditions in the exposed and cave deposits. This supports the inference that both the sediments and fossils are allochthonous. Thus, the red cave soils are being transported from the above scarp into the cavities. Mineralization of fossils within the deposit suggest mild diagenesis through infiltrating water. However, the isotope values yielded by the tested samples indicated little or no major diagenesis other than slight mineralization.

Deposition seems to have been slow as is suggested by the marked stratigraphic architecture and the slow sedimentation rates calculated for several of the intervals. Layer or bed architecture was variable, several layers were separated by discernable disconformities that mark different erosional/depositional events and changes in sedimentation regimes (Figure 3-4). The beds were generally prograding, with the lowest layers representing lower energy (horizontal) depositions, whereas the upper-level layers were more amalgamated and inclined, suggestive of slightly higher energy flooding events resulting in more pronounced rill erosion. Several beds showed evidence of slump erosion and truncation likely caused by rill erosion (Figure 3-4). The weathering levels observed in osseous remains rarely surpassed stage 2, which suggests that the pellets and their content were exposed for only 2 to 4 years before final deposition and diagenesis, where they decomposed exposed to the air, thus attracting insects. This is likely to have occurred in the primary pellet deposit in the upper cave levels, and much before transportation into the cone deposit below.

One of these events (layer F up to C), suggested a stratigraphic inversion, mixture with a slightly faster sedimentation rate of > 1.3 mm/yr^-1^. Together, layers F–C may constitute a flooding event in which older fossils were transported and deposited over younger deposits, as suggested by the ^14^C AMS date for layer F, E and D. Bioturbation also could have been a major source of reworking and stratigraphic inversion (Bosch and White, 2007; Patzkowsky and Holland, 2012). Although most exotic taxa occurred in the upper intervals, the anomalous presence of *Rattus* spp., *Mus musculus*, and *Passer domesticus* within the lower levels and the older ^14^C date in level II support either mixing of diachronous fauna or a stratigraphic inversion at level II (Table 4; unp. Data from dated *Antrozous* and *Boromys*, see Orihuela et al., forthcoming). Land crabs, rodents and island-shrews are known to excavate and burrow in the sediment and for scavenging (Andrews, 1990) which can result in the mixing of diachronous remains. However, bioturbation index was low at most intervals, between 0 and 1 (i.e., 1–4 % overall bioturbation), except for interval II, which had a bioturbation index of 2 (>15%).

Furthermore, the native rodent and *Nesophontes* tooth marks reported in the assemblage (Orihuela et al. 2016), and the occurrence of a fly pupa and traces of insect activity on several of the bone remains (Figure 9.3, 10.1, 10.2, and 10.5) suggest that pellets laid exposed long enough to attract these scavengers before final deposition in the cone deposit. Overall, this supports the mixing of fauna in the upper primary deposits, causing some of the events and specimens to reach the deposition cone already mixed, or being further mixed there.

The large accumulation of gastropods, ash, and charcoal detritus in layer C suggests another major deposition event. Bed C registers a probable large forest fire in the upper scarp and wooded areas above the cave. In general, the material from the major events indicated by beds C, E, and F, was very poorly sorted with well-preserved fossils, seeds, and plant material. This suggests that these layers may represent diamicton facies of Gillieson (1986), which could be interpreted as large asynchronous flooding events (McFarlane and Lundberg, 2007), although in a restricted smaller scale. In turn, the slow sedimentation rates, weathering levels, and fly pupae imply longer times of non-deposition, exposure, and erosion. The amalgamated mixture of larger and smaller vertebrates with land gastropods suggests that deposition is largely controlled by turbulent flooding events of moderate energy (Farrand, 2001; McFarlane and Lundberg, 2007). This is further supported by an observation. In April 2015, two of us (JO and LPO) experienced a torrential rainstorm under the main doline, but it failed to bring material into the deposit cone, suggesting that the transportation events must be of a more intense nature in order to transport sediment and biological remains into the cave. Interestingly, some of the superficial dates acquired for the upper levels (*n* = 3: 1953–1957 AD) agree with a period of prolonged rainfall and inundation in the region (Pérez et al., 2017).

### Taphonomy: raptors as one of the deposit-formation processes

Because these faunal remains are the results of raptor predation, they represent a fauna of regional or local scale, but not collected by a single raptor. *Tyto furcata*, the most common of Cuban nocturnal raptors today (Garrido and Kirkconnell, 2000), is a small mammal specialist with a hunting radius between 3 and ∼ 16 km (Banks, 1965; Andrews, 1990) and is probably one of the major contributors to pellet accumulations in Cuba today (Arredondo and Chirino, 2002; Silva et al. 2007; Hernández and Mancina, 2011, López, 2012) and the major contributor to the formation of the doline deposit.

*Tyto* species of barn owls were formerly considered a non-preferential predator (Bunn et al. 1982). Today they are regarded as highly selective (Andrews, 1990; Kusmer, 1990; Hernández and Mancina, 2011), with prey that range in weight between 25 and 200 g, but of which over 95 % of prey items weigh less than 100 g (Morris, 1979). Diet studies of *T. furcata* in Cuba show that bats, reptiles and birds constitute a small percentage (< 5 %) in the diet, whereas rodents, especially the exotic murids, make up more than half of their diet (Silva, 1979; Suárez, 1998; Arredondo and Chirino, 2002; Hernández and Mancina, 2011; Lopez, 2012). Among the bats, those species with stationary feeding habits, such as *A. jamaicensis*, *Brachyphylla nana,* and *Phyllonycteris poeyi*, are the most common species present in pellets (Silva, 1979; Hernández and Mancina, 2011; López, 2012). Other species with similar feeding such as *P. falcatus* and *Erophylla sezekorni* are also frequent (Silva, 1979; Arredondo and Chirino, 2002; Hernández and Mancina, 2011).

The high preference for exotic murids (*Mus* sp. and *Rattus* spp.) is likely a post-Columbian adaptation that replaced reliance on *Nesophontes*, bats, and birds in the past. Studies have shown that where rodents are not available, bats, lipotyphlans, and birds make up most of the diet (e.g. Velarde et al., 2007). This hypothesis can help explain their higher frequency in this and other Antillean deposits.

The most abundant fauna encountered in our assemblage range in body mass from 5g to ∼ 1000g (∼ 1 kg); from the smallest bats to the small-medium sized capromyid rodents such as the *Mesocapromys* spp., *Boromys* spp., and juvenile *Geocapromys columbianus* (all >160g or 0.16 kg), plus *Capromys pilorides* which is heavier (> 1 kg) (supplement in Turvey and Fritz, 2011; Borroto-Páez and Mancina, 2017).

Thus, it is likely that *T. furcata* was not the sole contributor to the pellet-derived fauna reported here, for there were more strigids in Cuba’s past, and at least three extinct *Tyto* species (Suárez and Olson, 2015; Orihuela, 2019). Indications of multiple species of raptors contributing pellets to the deposit are suggested by the taphonomic evidence. One is the dominance in the frequency of cranial elements (skulls and mandibles) over long bones and other elements of the appendicular skeleton. This ratio is common in *Tyto*-derived pellet deposits but also in those of strigids (Andrews, 1990; Kusmer, 1990). In this assemblage, cranial elements were represented by 476 skulls (mostly incomplete with clear evidence of predation >45 %) and 1359 dentaries, constituting over 46 % of the total (or 1835 of total 3932) and 78% of the NISP. In comparison, non-cranial elements represented 17.3% of the NISP and 10.2% of the total remains. But their lower count is likely a bias of the collection effort, as the diversity indices, discussed ahead, imply. Overall, the assemblage had well preserved post-cranial elements, but until the full study is resumed we cannot determine whether post-cranial elements were more frequent than cranial elements, which would be suggestive of other medium-sized nocturnal raptors (Andrews, 1990).

Evidence that more than one raptor species was involved in the deposition of pellets is further suggested by the increased diversity and presence of predominantly larger fauna, and by the partially mineralized large pellets found within beds G and H. These two layers were especially rich in juvenile capromyid rodents (> 400 g), larger birds and bats (Table 3). These raptors could include other Cuban extinct tytonids or strigids, such as larger and diverse *Tyto* (e.g., *cravesae* or *noeli*) taxa or *Pulsatrix arredondoi,* based on the diversity of the faunal assemblage (e.g., see Restrepo-Cardona et al., 2018). Of these, Arredondo’s spectacled owl *P. arredondoi* has been confirmed to have survived into the very Late Holocene (Jiménez et al., in press), which can also be the case for other Cuban extinct raptors (Orihuela, 2019), and at this point it cannot be excluded as contributor to the deposit formation. Pellet studies of *P. perspicillata* showed a wide diversity in avian prey items, including hummingbirds and migratory species (Restrepo-Cardona et al. 2018).

Extant strigids cannot also be ruled out. These may include *Asio*, *Otus*, or *Margarobyas lawrencii* and *Glaucidium siju*, several of which inhabited and still inhabit the region (Jiménez, 1997, 2001; Jiménez and Arrazcaeta, 2015; Garrido and Kirkconnell, 2000). The remains of some of these owls are present in our assemblage, likely as a result also of raptor predation. *Tyto furcata* remains were also found; all either as results of raptor predation or natural death.

Thus, it is likely that more than one predator, either extant or extinct, contributed to Cueva de los Nesofontes’s doline deposit over the span of 2000 years. A multi-raptor deposit in the same cave has already been suggested (Orihuela, 2010). Generally, raptor pellets provide a good record of local or regional fauna because of their broad spectrum of selectivity of available microfauna (Mikkola, 1983; Andrew, 1990; Kusmer, 1990). In Cuba, pellet studies have shown that such selectivity does not vary significantly across habitats, whether disturbed or natural (Hernández and Mancina, 2011). A larger source for bone accumulations that include both natural and raptor-derived faunas widens the diversity of our record. In that sense, our subsample could be a good proxy of a past local or regional land vertebrate fauna.

### Fauna Diversity

The faunas recovered from this assemblage are moderate to highly diverse (83 NTAXA; 73 vertebrates) and somewhat homogeneous (Shannon-Weiner index of 2.82). Among the vertebrates, the relative abundance was particularly highest in birds and mammals (Simpson dominance >0.293 or 29.3%), of which *Nesophontes* and *Artibeus* spp. made up the largest NISP (Table 6). Individually per stratigraphic interval, the homogeneity index (Shannon-Weiner) and evenness index varied between 1.17 and 1.21, and 0.81 and 0.83 between interval levels, respectively. The highest being level IV (1.21; 0.83), and the lowest level II (1.17; 0.81) (Table 6). However, there was a poor negative (linear) correlation (R²=0.395) between the Shannon-Weiner index and NTAXA. These suggested, nevertheless, that the stratigraphically lower and chronologically oldest intervals II and IV were less diverse, whereas the youngest I and III were more diverse and thus less homogeneous, but better representatives of the collective fauna. The Fisher ά and Simpson’s indices reflect the higher diversity of levels II and IV (Table 6; Figure 12). In this sense, heterogeneity could have been a result of sample recovery variation, overall and between intervals, and the fauna diversity present therein. The NISP of our assemblage nearly reached an NTAXA asymptote after 3000 specimens and over 70 taxa, suggesting that our overall sample size approached maximum diversity in vertebrates expected for the deposit, but not so each individual bed (Figure 11).

**Figure 11:**
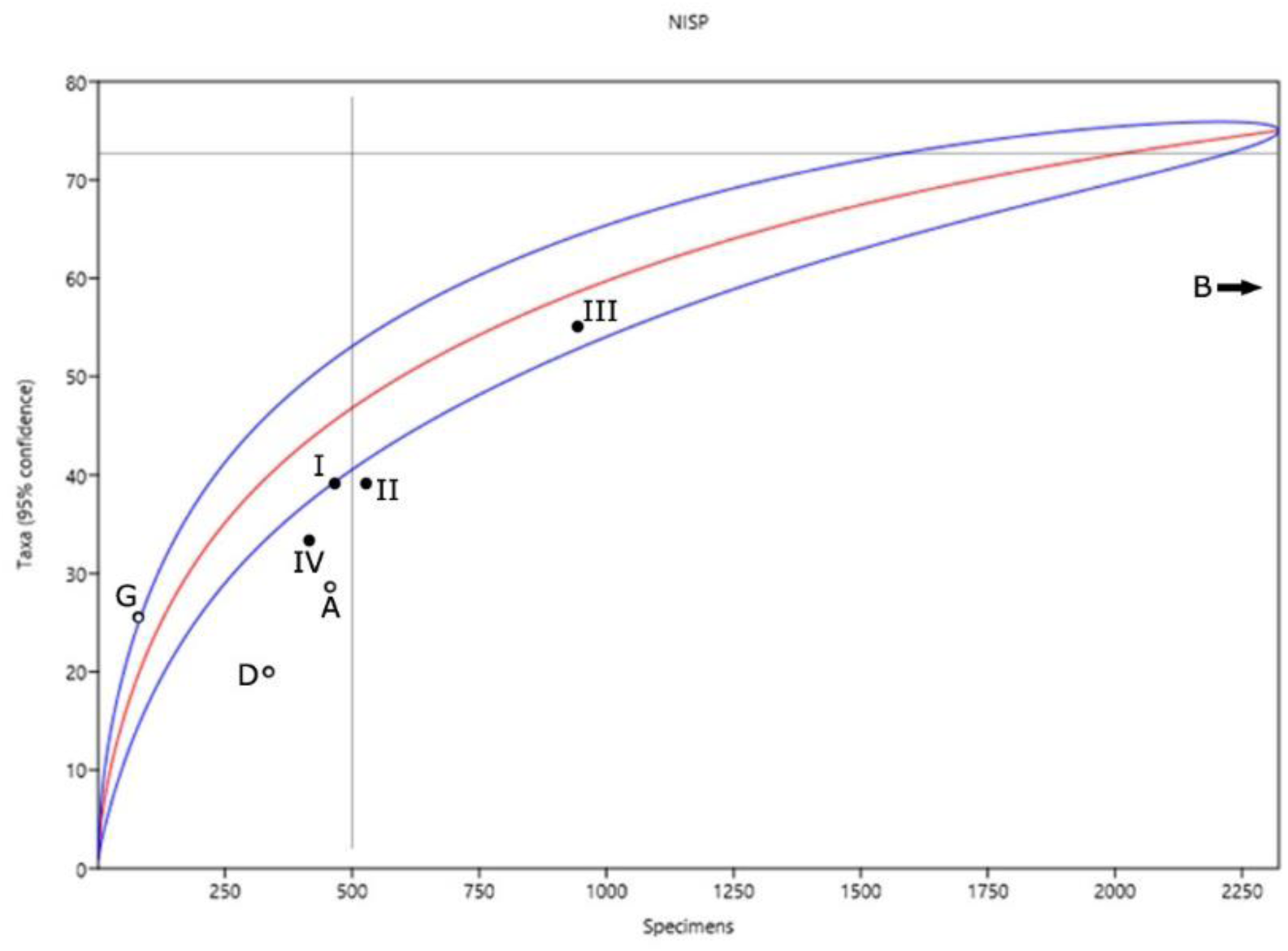
Rarefaction curve of Cueva de los Nesofontes doline test pit D (Levels I-IV) in relation to other Cuban deposits (A-G). A is Cueva GEDA, Pinar del Río. B is Cuevas Blancas, Mayabeque. D is the *Desmodus* deposit described in Orihuela (2010). G is from Gato Jíbaro archaeological deposit described in Orihuela and Tejedor (2012).

**Figure 12:**
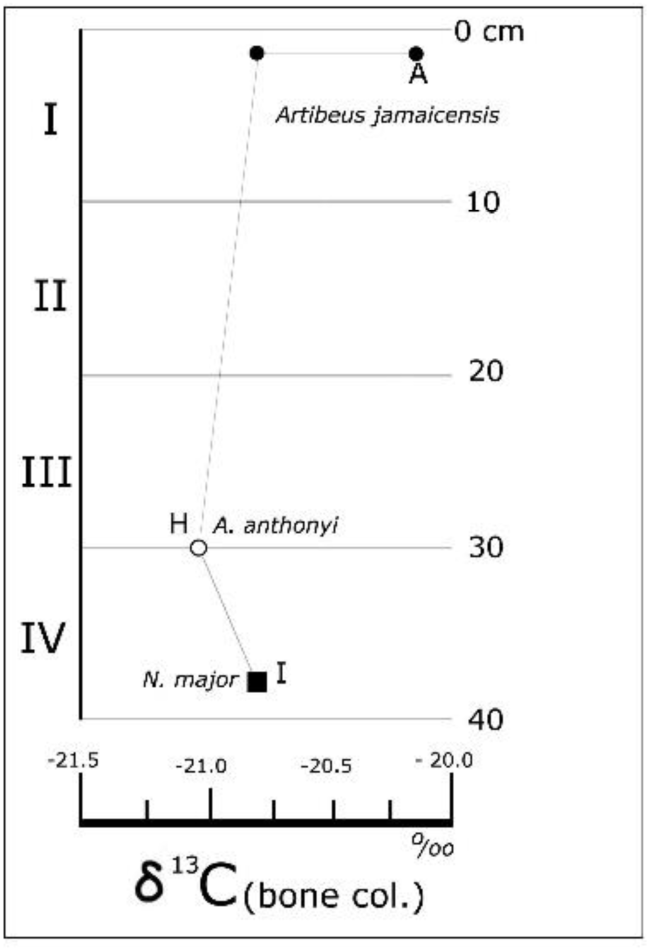
Carbon stable isotopes signals from *Nesophontes* and *Artibeus* spp. (from bone collagen).

**Table 6:** Diversity indices, evenness, dominance, and chronology for each stratigraphic interval. Overall values are not the averages of each column, but the overall index calculated for the whole fauna. R = recent.

| Interval | NTAXA | NISP | Shannon-Weiner | Simpson's | Fisher-a | Eveness | Dominance | Chronology |
| --- | --- | --- | --- | --- | --- | --- | --- | --- |
| I | 39 | 469 | 1.18 | 0.653 | 1.12 | 0.814 | 0.525 | AD 1953-1993 |
| II | 38 | 519 | 1.17 | 0.649 | 1.14 | 0.805 | 0.572 | AD 660-770 |
| III | 55 | 929 | 1.18 | 0.653 | 0.99 | 0.812 | 0.481 | AD 605-655 |
| IV | 34 | 409 | 1.21 | 0.664 | 1.18 | 0.839 | 0.545 | BC 40-90 AD |
| Overall | 73 | 2326 | 2.82 | 0.698 | 1.1075 | 0.233 | 0.53075 | R-2000 BP |

A preliminary comparison in NTAXA and NISP between several of Cuba’s cave accumulation deposits can help further contrast the richness of the main doline deposit in Cueva de los Nesofontes and characterize the diversity of the assemblage. Even so, any comparison of homogeneity, diversity and evenness indices among Cuban cave deposits is limited due to the lack of comparable published assemblage details for other caves that allow for such calculations, and because the deposits have a different genesis and were sampled or studied differently (e.g., concentrated on different groups of organisms, as may seem obvious). For example, the assemblages reported with appropriate detail for Cueva de los Masones and Jagüey, in Sancti Spíritus, represents bats (Silva, 1974), whereas other such as Cueva GEDA in Pinar del Río (Mancina and García-Rivera, 2005; Condis unp. Thesis), and other cave deposits in northwestern Cuba (Orihuela, 2010; Orihuela and Tejedor, 2012) included several groups of vertebrates, but were less diverse in NTAXA (between 20 and 29) with smaller sample collections (between 150 and 430 NISP). Stratigraphic details on NTAXA and NISP variation from other faunistically-rich cave deposits such as Cueva del Túnel, Cueva del Mono Fósil or Cueva de los Paredones are not available.

The rarefaction asymptote for Cuevas Blancas (Mayabeque), Masones and Jagüey deposits extend well beyond the 500 NISP but with lower NTAXA than the deposit reported here. Cueva GEDAS (Condis, unp. Thesis), the kitchen midden from Cueva del Gato Jíbaro (Matanzas; Orihuela and Tejedor, 2012; JO unp. data), and the other deposit reported for Cueva de los Nesofontes (Orihuela, 2010) all cluster behind the 500 NISP level, and lie outside the confidence intervals, which suggests under-sampling (Figure 11).

Our deposit was richer than that of Cuevas Blancas in NTAXA vertebrate diversity (*n* = 83 vs. 59), even though this last had a much larger NISP sample size (i.e., 10,027 vs. 2326) (Jiménez et al., 2005) (compare to B in Figure 11). Only the midden deposit from Gato Jíbaro and intervals I and III of Cueva de los Nesofontes doline deposit reported here were within the confidence interval of the curve (Figure 11). The remaining assemblages were outside and were less than the 500 NISP mark, also suggesting under-sampling.

The assemblage’s diversity has been influenced by our sampling methods and differences in taphonomic aspects such as raptor preference and natural death, but also by reworking and sedimentological processes explained above. In paleontology, one can never have full access to the actual original faunal diversity. But in this sense, our calculations allowed us to quantify diversity and compare it to other important Cuban deposits to see where our assemblage fits and explore what that says about its diversity and formational history. Calculating the diversity indices for each bed and the assemblage of Pit D, has permitted us to compare the diversities of each subsample (each bed), and to infer that the high diversity observed is partially representative of the local fauna, despite limited sampling and completeness, due to the multiple origins of the biological remains. Thus, suggesting that the presence of several groups or taxa are more than taphonomically or raptor selection, but also controlled by the sedimentological history of the deposit. And moreover, that this is one of the most diverse paleontological cave deposits studied from Cuba, and its further study can provide a noteworthy contribution to Cuban and Antillean vertebrate paleontology.

### Chronology and fauna contemporaneity

Since the transport and deposition occurred after the deposition of pellets in the upper levels (i.e., primary deposit), the fossils transported and incorporated in the doline deposit below act as *terminus post quem* (TPQ) to the formation of the beds. Therefore, the depositional/erosional events indicated by the disconformities should date to a time after the age of death of the fossils. In this sense, beds I and H and B and A seem to follow the law of deposition, whereas the events recorded in layers C-F, may have occurred after the deposition of the layers H and G, incorporating an older non-contemporaneous faunal assemblage (Figure 3– 4). This is important in the interpretation of the contemporaneity and diachrony of the fauna contained on each bed as a depositional event.

The problem of contemporaneity in fossil assemblages can be further complicated by bioturbation, mixing, layer inversion, down-slope truncation, and further reworking (Farrand, 2001; Bosch and White, 2007; McFarlane and Lundberg, 2007; Patzkowsky and Holland, 2012). Although it is often not practical due to cost or preservation, it is preferable that many specimens from a single layer or a whole stratigraphic sequence are dated (e.g., see Semken et al., 2010; Stoetzel et al., 2016). Several studies have suggested that direct dating of associated specimens is required to establish whether specimens in the same bed are radiometrically contemporaneous or diachronous (Stafford et al., 1999; Stoetzel et al., 2016). Semken and colleagues showed that diachrony is generally the norm, as they shown in several North American deposits (Semken et al., 2010), this may be the case in Cuban cave deposits as well.

These issues are of great concern in the study of Cuban bone accumulation assemblages, the majority which today lack ^14^C dates. When available, they are often single dates that do not follow a stratigraphic sequence or form a suite, and, as we have encountered here, may represent diachronous faunas. Thus, assessing contemporaneity between important assemblages and their LADs remains a key factor in the study of extinct or extirpated faunas, but is largely unachievable in Cuba until more dates are available. This is a hindrance to the understanding of Cuban, and thus Greater Antillean, vertebrate extinction and faunal turnover since the late Pleistocene and through the Holocene. A comparison of faunas is further augmented by the lack of confirmed late Pleistocene dated deposits in Cuba. Several candidate faunas have been postulated (e.g., Cueva del Mono Fósil, Cueva del Túnel or Cueva de los Paredones; Salgado et al., 1992; Gutiérrez et al., 2014; see Figure 1), but only three (Iturralde-Vinent et al., 2000, Breas de San Felipe, El Abrón, and Ciego Montero) have been confirmed (Kulp, 1952, Suárez and Díaz-Franco, 2003; Jull et al., 2004; Fiol, 2015). Conversely, many specimens from deposits that were originally thought to be at least late Pleistocene in age have yielded much more recent dates (mHOL-lHOL; MacPhee et al., 1999, 2007; Jiménez et al., 2005; Jull et al., 2004; Orihuela, 2010; Orihuela, 2019; Orihuela et al., forthcoming).

In our assemblage, the faunas of bed G and H (level intervals III-IV) can be interpreted as near contemporaneous, since they differ in slightly less than two sigmas (2σ: 100–140 ^14^C cal. deviation years). A deviation of a single sigma, usually between 60–70 ^14^C cal. years is preferable (Semken et al., 2010), but not available for this deposit. But overall, our dated intervals (e.g., F, E, C and B) are generally longer than 1σ or 2σ, and cannot be considered fully contemporaneous. The difference in diachronic range is between 118 and 138 cal. yrs. BP, among the faunas of intervals III and IV, and of ∼1958 years between I and II. The diachrony between these intervals highlights wide temporal hiatuses that support a non-continuous deposition, and likely, asynchronous faunas above bed G due to the processes already discussed.

The use of a single date, even if from a single important individual extracted from a controlled stratigraphic unit, can conflict with or non-representative of the age of the whole fauna present in a unit or its stratigraphic association, as is suggested by the ^14^C age of *P. vetus* from layer E. As is the case in many studies of Antillean land vertebrate paleontology, the interpretation of a single date as a representative of unit-fauna contemporaneity must be considered cautiously. Our data support the use of multiple dates, acquired directly from identifiable bone specimens, in the study of assemblage faunas. Better yet, several specimens should be dated within the same stratigraphic unit, or whole stratigraphic suites when possible in order to understand depositional regimes, spatial-temporal faunal change, diachrony, and bio-ecological turnover.

However, we consider that even though our dated individuals are not strictly contemporaneous, (as they are not expected to be in a time-averaged, slowly formed deposit), the direct LADs they provide for extinct and extirpated taxa are useful to biogeographical discussions (MacPhee et al., 1999; Silva et al., 2007; Patzkowsky and Holland, 2012). All direct ^14^C dates provided for the extinct fruit bats *A. anthonyi* and *P. vetus* and the island island-shrews *Nesophontes* spp. provide evidence of their survival/existence, well into the very late Holocene of Cuba. The chronological and stratigraphic evidence suggests that the studied deposits are about 2000 years old, at least to the level excavated and thus includes fauna from well within the pre-Columbian Amerindian interval (Morgan and Woods, 1986; Cooke et al., 2017).

Furthermore, this indicates not only post-Pleistocene-early Holocene survivorship but also wider distribution ranges that persisted for several thousands of years of climate variations and human coexistence into the Late Holocene. Further supporting the time-lagged, group-specific asynchronous extinctions hypothesized by MacPhee and colleagues (1999), which have received growing support in Cuba (Jiménez et al., 2005; Steadman et al., 2005; Orihuela, 2010, 2019; Orihuela and Tejedor, 2012; Borroto-Páez and Mancina, 2017; Orihuela et al., forthcoming).

### Fauna temporal-spatial distribution

Several of our fauna records indicate a wider distribution beyond current limits for several species that lasted well after 2000 years BP. These include the anole lizard *Anolis* cf. *chamaeleonides*, the woodpecker *Colaptes fernandinae* (or *auratus*), the Cuban parakeet *Psittacara eups*, and crow *Corvus* which are today locally extinct in the region surrounding Palenque. These represent past extralimital records for species whose distributions lie currently far from the deposit (Garrido and Kirkconnell, 2000; Rodríguez-Schettino et al., 2013; Orihuela, 2013). Other remains constitute the first pre-Columbian, paleontological records for *Progne* cf. *subis*, *Tachycineta bicolor* and *Cathartes aura*. *Cathartes aura*, *Corvus* sp, and *Psittacara eups* have been reported from colonial contexts of the 16th and 18th centuries of La Habana Vieja (Old Havana) (Jiménez and Arrazcaeta, 2008, 2015). The extralimital presence of *Corvus* sp. in the colonial contexts of the old city of La Habana seems to support a recent range constriction likely related to deforestation (Jiménez and Arrazcaeta, 2008; Orihuela, 2013). *Cathartes aura* was initially reported from a supposed late Pleistocene deposit of Cueva del Túnel, in Mayabeque province (Acevedo et al., 1975; Acevedo and Arredondo, 1982). That report was challenged by Suárez (2001), who identified those specimen as modern (Jiménez and Arrazcaeta, 2008), thus deleting the species from the fossil record of Cuba. Moreover, Suárez (2001) indicated the existence of an undescribed species of *Cathartes*. The turkey vulture was observed and sketched by a British soldier during the siege of Havana city in the summer of 1762 (campaign journal of Henry Fletcher, 1757–1765: 255).

All of the rodent species had already been reported for the region and do not constitute new records (Jiménez et al., 2005; Silva et al., 2007; Orihuela and Tejedor, 2012). The *Mesocapromys nanus* and *M*. *kraglievichi* are interesting because their fossils support a wider late Holocene distributional range and several thousand-year survival post-Pleistocene climate change and human inhabitancy in the island. The survival of the extinct hutia *M. kraglievichi* through the pre-Columbian (Amerindian) interval is validated by the direct ^14^C LAD obtained from a specimen from the Solapa del Megalocnus site, Mayabeque province (Jiménez and JO unp. data). This specimen yielded an age of 1780±50 rcyr BP from one of the preceramic archaeological contexts, but the specimens found between intervals III and IV of Cueva de los Nesofontes doline assemblage suggest a slightly younger LAD for this species. *M. nanus* is today likely extinct, formerly restricted only to the Zapata swamp, but in the past, it had a wider range (Silva et al., 2007; Borroto-Páez and Mancina, 2017). A similar extralimital fossil record was recently provided for *Mesocapromys sanfelipensis* on the mainland of Cuba (Viñola et al., 2018). This taxon is one of three highly localized and endangered pygmy hutias found today exclusively on several keys of the Cuban archipelago (Borroto-Páez, 2011; Mancina, 2012).

Bats and *Nesophontes* were the most abundant vertebrates in the assemblage. Yet, several of their species were rare and appeared only at specific intervals or beds, such as *Nesophontes* cf. *longirostris*. The smaller *N. micrus* dominated this genus’ frequency, with more than 600 NISP representing at least 62 individuals (MNI) present at all intervals. But, individuals of *N. major* were slightly more abundant (Table 3–4). Of all *Nesophontes* species, *N*. cf. *longirostris* was the scarcest, further supporting the rarity of this species (Anthony, 1919). Over 2000 near-complete crania of *Nesophontes* spp. were formerly extracted from the 1985 excavation alone, making this one of the richest *Nesophontes* bone accumulations reported from Cuba (Vento, 1985 in Nuñez, 1990, vol. 1: 299–304).

The bats were especially numerous and diverse. The 18 taxa recorded here represent more than half of the known Cuban bat fauna. The taxonomic diversity of bats increases to 21 species if other species documented for this cave are counted (i.e., *Desmodus rotundus*, *Lasiurus insularis* and *Chilonatalus macer* in Orihuela, 2010).

The assemblage was particularly rich in frugivorous bats *B. nana*, *Phyllonycteris poeyi*, *jamaicensis,* and *P. falcatus*, whereas the *insectivorous* bats *Eptesicus fuscus*, *Tadarida brasiliensis*, *Mormoops blainvillei, Pteronotus parnelli,* and *Macrotus waterhouseii* were less common (Table 3). This could be a result of predator selection as aforementioned. The predators that contributed to the pellet-derived deposit seem to have targeted small or medium-sized gregarious species with stationary feeding habits (phyllostomids) or species that had accessible roosts (molossids). Aerial, fast-flying insectivores such as the molossids, and the larger fish and blood-feeders (e.g., *Noctilio* and *Desmodus*) hardly ever occur in owl pellet deposits, which in part can explain their rarity in Cuba’s raptor-derived bone deposits.

*Molossus molossus* was represented in our assemblage by three partial skulls with evidence of predation and digestion found only in the uppermost two layers (A–B) of the first interval (Level I). The rarity of *M. molossus* in our assemblage can be the result of a more recent adaptation of both *M. molossus* and raptors such as *Tyto furcata*. *Molossus* species are rare in the Cuban Quaternary cave deposits likely because before European arrival these species roosted in trees or crevices, which are not prone to intense preservation.

### Coexistence and competition

Specimens of *Artibeus anthonyi* and *Artibeus jamaicensis* occurred in direct association within the same layer unit and throughout two intervals (level III and IV). *Phyllops vetus* and *Phyllops falcatus* occurred together only at interval level III (beds G and H). Interestingly, in bed E of interval II, which yielded the direct date for *P. vetus* and the oldest ^14^C available for the assemblage, the two *Phyllops* species did not coincide. In the youngest interval (level I), neither the extinct *A. anthonyi* or *P. vetus* occurred, suggesting that by then they were not predated by raptors, were rare to appear in the record, or already extinct. Nonetheless, this supports a very Late Holocene extinction for these two species. Moreover, this record suggests that today’s most endangered and rarest of Cuban bats, *Natalus primus* and *Antrozous koopmani*, had much better distribution in the island that lasted up to very recently.

The direct ^14^C date on *A. anthonyi* reported for this interval suggests that it is highly probable that both *Artibeus* species coexisted for several thousand years. It was recently considered that these taxa did not coexist in the Holocene of Cuba for lack of direct evidence (Balseiro et al., 2009; Turvey and Fritz, 2011). In a few deposits where *A. anthonyi* occurred, *A. jamaicensis* was not found, and when found, the fossils seemed to be non-contemporaneous. But this has not been the case in others. H. E. Anthony, in the original description of the specimens from Cueva del Indio in Daiquirí, Eastern Cuba, that was later identified as *Artibeus anthonyi* by Woloszyn and Silva (1976), mentioned the occurrence of both *Artibeus* species, but these specimens have not been dated.

Until now, a direct radiocarbon date on *Artibeus anthonyi* was unavailable, but other evidence already suggested coexistence and survival well into the Holocene of Cuba (Jiménez et al., 2005; Orihuela, 2010; Condis unp. thesis). Recently, Condis reconsidered the temporal coexistence of several of Cuba’s extinct bats in Cueva GEDA, including *Cubanycteris silvai*, *P. vetus*, *A. jamaicensis* and *A. anthonyi* (Condis unp. Thesis). *A. anthonyi*, *P. vetus*, *M. megalophylla,* and *A. koopmani* have been reported in deposits dated between the Late Pleistocene (21,474–20,050 BP) of Cueva El Abrón (Suárez and Díaz-Franco, 2003; Fiol, 2015), the early-mid Holocene of Cuevas Blancas (7044–6504 BP) in Jiménez et al. (2005) or further in the very Late Holocene (Orihuela, 2010; Orihuela and Tejedor, 2012; Orihuela et al., forthcoming). This supports their somewhat continuous presence in the fauna since the LGM and throughout most of the Holocene up to the colonial period.

Woloszyn and Silva (1977) suggested that the extinction of *A. anthonyi* was caused by competition, as may also be the case for *P. vetus* and *P. silvai*. However, this hypothesis lacks confirmatory evidence (Balseiro et al., 2009). With the stable isotope values acquired from the bone collagen and tooth apatite from *A. jamaicensis* in comparison to *A. anthonyi* specimens, we are in a better position to discuss the competition hypothesis. The carbon isotopes do not indicate a substantial trophic variance between the species (Table 2; Figure 12). *Artibeus anthonyi* and *A. jamaicensis* had a similar diet and occupied a similar niche, as suggested by their values: *A. anthonyi* (δ¹³C_col. −21.1 ‰ and δ¹³C_apt. −11.0 ‰) and *A. jamaicensis* (δ¹³C_col. −20.1 and −20.7 ‰, plus δ¹³C_apt. −8.1 and −9.9 ‰). These values suggest that the component of diet could have been an important source of competition; the intensity of the competition depending on their level of resource partitioning or difference in foraging strategies is yet unknown, and only here incipiently investigated and requiring further data.

Moreover, our *Artibeus* δ¹³C_col. values were lower than those reported by Rex and colleagues for *A. jamaicensis* (−25.6±0.55 SD, *n* = 17) and *A. lituratus* (−25.2±0.46 SD, *n* = 29) (Rex et al., 2011, p. 221). The slightly smaller isotopic yield of *A. lituratus* could suggest a slight vertical stratification in niche partitioning between these two species in Neotropical forests, following the hypothesis that smaller bats prefer understory resources, whereas larger species prefer larger fruits of the canopy (Findley, 1993; Bonaccorso et al., 2007; Pereira et al., 2010). *A. jamaicensis* generally feed in the forest understory, commonly at ground level, whereas *A. lituratus* preferred a higher canopy level (McNab, 1971; Herrera et al., 2001; Rex et al., 2011; Silva et al., 2008). However, this was not supported overall for phyllostomids (Rex et al. (2011). Rex and colleagues reported that there could be up to ∼ 6.8 ‰ carbon isotope variation between syntopic species and concluded that there was no vertical stratification for these and other South American forest phyllostomids (Rex et al., 2011). This 6.8 ‰ value is much greater than the one we report for *A. jamaicensis* and *A. anthonyi* (Table 2, Figure 12). Thus, it is likely that syntopic phyllostomids such as *A. jamaicensis* and *A. anthonyi* explored all vertical forest levels and niches. The same phenomenon could have occurred among other extinct phyllostomids in Cuba. The size difference between *A. jamaicensis* and *A. anthonyi* is not considerable, and both can be classified as large short-faced fruit consumers (Silva, 1979). Therefore, their sympatry and minor habitat partitioning probably lead to more competition in foraging for the same resources in the same habitats.

Sympatry between the extant syntopic Cuban mormoopids and other Antillean nectarivorous phyllostomids, in which species presented differences in feeding apparatus, wing morphology, flight patterns, foraging behavior, and spatial segregation, probably facilitated resource partitioning (McNab, 1971; Herrera et al., 2001; Mancina and Herrera, 2010; Mancina et al., 2012; Soto-Centeno et al., 2014), and thus could have decreased competition.

Based on our few isotope values we hypothesize that *A. jamaicensis* and *A. anthonyi* had a similar diet and occupied similar habitats. In that sense, it is probable that *A. anthonyi* and *A. jamaicensis*, as for *Phyllops* spp., shared similar habitats and diets, because differences between the carbon and oxygen isotope values are likely not reflective of significant spatial segregation or foraging strategy. A higher level of competition in foraging habitats between these taxa could have pushed the rarer *A. anthonyi* and *P. vetus/silvai* to become extinct. A similar situation could help explain the coexistence of several extinct Cuban bats for a few thousand years and provide a window into their interaction and extinction We consider *A. anthonyi* rarer because in the cases in which both species occur, *A. jamaicensis* is by far the most common species of the two, as is the case in assemblages from Cueva de los Nesofontes (Orihuela, 2010; this work), Cueva GEDA (Balseiro et al., 2009; Condis, unp. Thesis.) and Cuevas Blancas (Jiménez et al., 2005). Although, this apparent variation in abundance could be a taphonomic artifact, such as raptor preference, and not reflective of their natural abundance. However, further isotopic analyses are required to corroborate these preliminary observations.

### Paleoenvironmental reconstruction gleaned from fauna and isotopes

The presence of the birds *Sturnella magna*, *Melanerpes superciliaris*, *Corvus* sp. and *Colaptes* woodpeckers suggest the presence of savannas and grasslands and nearby dry semideciduous forests. These are habitats which are today reduced, but still available over the karst terrain. Similar findings were reported for Cuevas Blancas, several dozen kilometers to the southwest of Palenque (Jiménez et al., 2005). The dove *Geotrygon chrysia* suggests dry forests with little undergrowth, and *Psittacara eups* undisturbed forests and palm grove savannas (Garrido and Kirkconnell, 2000). The regular transient woodpeckers *Sphyrapicus varius*, swallow *Tachycineta* sp. and the martin *Progn*e sp. support the presence of seasonal transient species in the assemblage. This mosaic of available habitats agrees with the former vegetation hypothesized for the region (Marrero, 1972; Del Risco, 1989). The pollen, spores, and seeds registered from this deposit are yet to be studied but can provide a better record of vegetation change in the area when available (Figure 9).

The carbon and oxygen isotopes aid in the interpretation of past habitats and microenvironments (Bocherens et al., 1996; Lee-Thorp et al., 1989; MacFadden et al., 1996). Habitats with a greater proportion of C4 vegetation, such as grasslands and savannahs, generally yield higher δ^13^C (more positive) values, whereas habitats with higher tree cover (riverine woodlands) tend to have lower (more negative) values. Mixed habitats yield intermediate values (Leichliter et al. 2016; Keicher et al., 2017). The δ^13^C values we obtained from the analyzed remains of *Artibeus* sp. and *N. major* are intermediate (Figure 11), suggest that these species lived in riverine woodlands and mixed woodland habitats. In this sense, the slight δ^13^C variation between the Cuban *Artibeus* spp. (δ^13^C_col. −21.1 and −20.1 ‰) discussed above suggests that *A. jamaicensis* and *A. anthonyi* probably preferred similar forest microhabitats within mixed woodlands and riverine woodlands.

Apparently, *N. major* inhabited similar habitats. Based on δ^13^C_col. values reported for *N. micrus* by MacPhee et al (1999 p. 16), which varied between −18.9 and −19.7 (*n* = 2), we infer that there might have been microhabitat segregation or resource partitioning (slightly different dietary niches) between the Cuban *Nesophontes* species, with *N. major* preferring mixed woodlands with more tree cover and *N. micrus* preferring grasslands and savannahs. If confirmed by further tests, this observation could explain the higher frequency of *N. micrus* relative to *N. major* observed in our assemblage. Even though *N. micrus* is smaller, it would have been easier to capture by nocturnal raptors, such as *T. furcata*, which prefer to hunt in more open terrain (Andrews, 1990; López, 2012).

In terms of diet, the single acquired nitrogen and carbon isotope value suggest that individual fed on millipedes, earthworms, maybe fungi and fruits (see similar interpreted signals in Reid et al., 2013; Eckrich et al., 2018). Based on this we hypothesize that *Nesophontes* species were probably omnivores, occasionally feeding on beetles or millipedes attracted to decomposing pellets accumulated at the raptor roosts. Their tooth marks have been identified on the bones present on owl pellet biological remains (Orihuela et al., 2016; this paper). However, the carrion signal acquired from scavenging is not clear in our isotope data. Our isotope signals for *Nesophontes* could be masked by enrichment from feeding on necrophagic arthropods (Hocking et al., 2007).

Once more, many more analyses are needed to explore habitat segregation and diet of these vertebrates. Thus, is probable that niche overlap could have also existed between these sympatric taxa. Other sources of variation could include metabolic and isotope fractionation differences (enrichment fluctuations) among taxa, body mass, trophic level, and individual habitat preferences, as has been shown for soricid shrews (Baugh et al., 2004; Keicher et al., 2017) could further mask the isotope signals. Nevertheless, the isotopes provide an additional, here incipiently explored, source of insight that can explain *Nesophontes* spp. habitat preference, diet, and competition to better explain their extinction.

The presence of mormoopid and vampire bats suggests an overall warm climate during the time of deposition since these species do not inhabit boreal regions and their distribution in the Neotropics is limited by temperature (Vaughan and Bateman, 1970; McNab, 1973; Bonaccorso et al., 1992). This is supported by the oxygen stable isotopes acquired from bone hydroxyapatite and collagen from remains of the bats *A. jamaicensis*, *A. anthonyi* and from *N. major* in several intervals of the deposit (Table 2). Our record suggests a wetter, warmer climate around BC 40 – 90 AD and AD 605–655, which agrees with the large, slowly deposited but amalgamated bed sets of interval IV and III (beds I to H), expressive of large flooding depositional events.

A −0.4‰ positive oxygen isotope excursion occurred at AD 660–770, suggestive of drier, colder local conditions, with a subsequent progressive return to warmer, wetter conditions after and up to the present (Figure 13). This profile is comparable to conditions gleaned from the Late Pleistocene speleothem record of Río Secreto, Yucatan (Mexico) between 23 and 23.5 ka (Medina et al., 2017) and the temperature deviations (anomalies in Celsius degrees from modern temperature) in the order of −0.6 to −0.2 in Moberg et al. (2005) and Abrantes et al. (2017). The wetter, warmer period that followed also agrees with the data presented by these researchers.

**Figure 13:**
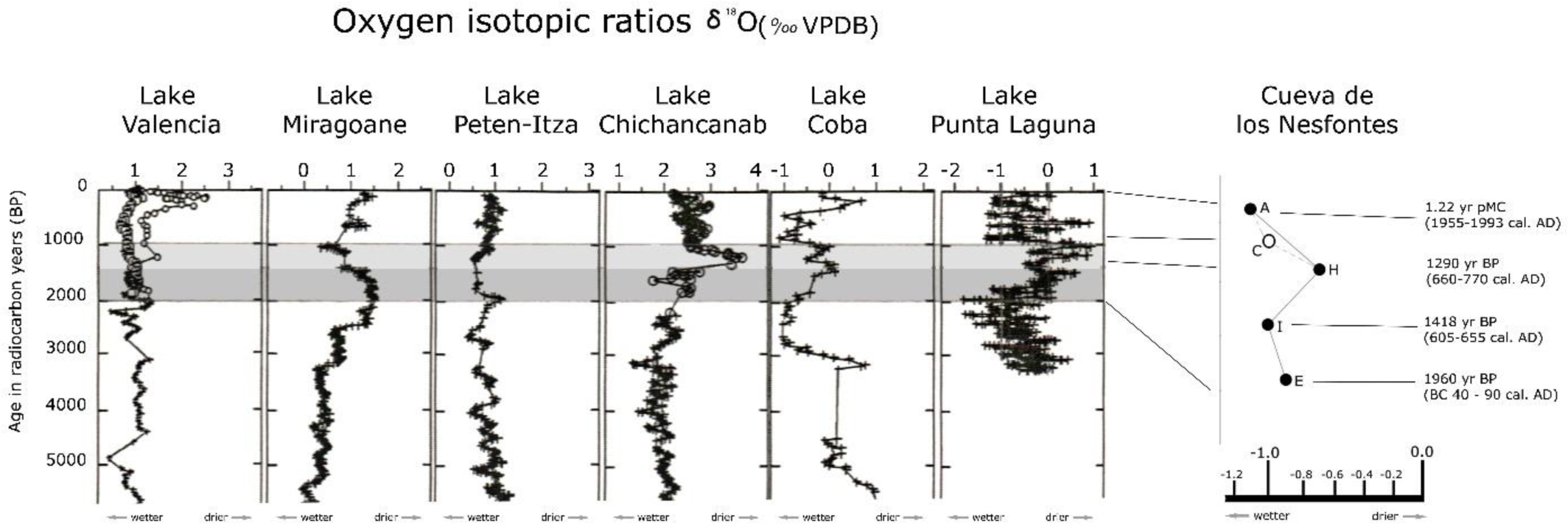
Approximation of paleoenvironment conditions through oxygen stable isotopes from Cueva de los Nesofontes test pit D, compared to other circum Caribbean deposits (modified from Curtis et al., 2001). The grey areas indicate the timeframe of our deposit and its graphical correlation to our data.

A similar positive excursion, although larger in magnitude, is recorded from lacustrine deposits from Punta Laguna, Lake Chichancanab and Lake Coba (Hodell et al., 1995; Higuera-Gundy et al., 1999; Curtis et al., 2001, p. 4). These records indicated a major drought between 1500 and 1100 years BP (op. cit.), which can help interpret our positive excursion as a similar, concomitant dry-cold spell.

Paleoclimatic records throughout the Caribbean and circum-Caribbean show a shift from wetter, more mesic conditions during the early-middle Holocene to drier, more xeric conditions during the late Holocene, between 3000 and 1300 year BP (Hodell et al., 1995; Curtis et al., 2001; Peros et al., 2007). Our data do not agree with the overall tendency towards drier conditions after 2000 cal yr. BP interpreted from Cuban coastal lacustrine deposits (Peros et al., 2007; Peros et al., 2015; Gregory et al., 2015), and instead agree with those acquired from cave speleothem records (Pajón et al., 2001; Pajón, 2012; Fensterer et al., 2013) which indicate the inverse. Since our oxygen isotope values were acquired from bat dental apatite, they likely represent the bat’s life oxygen record acquired from the local diet and water source (Bocherens et al., 1996; Lee-Thorp et al., 1989; MacFadden et al., 1996). These, in turn, provide us with a very local environmental record. Nevertheless, our contrasting results could also be obscured by the complicated fractionation of oxygen in the sampled bats, bone mineralization, or deposit diagenesis (Bocherens et al., 1996; Lachniet, 2009).

The large accumulation of charcoal and ash of bed C suggest either a natural forest fire or anthropogenic activity in the area; with the charcoal remains brought in by a fast flooding event. Direct ^14^C date estimates from this level (bed C in Figure 4) indicates it includes fauna younger than ∼ 1960 cal. yrs. BP (BC 40 – 90 AD), but older than 1000 BP (from *Boromys torrei* and *Antrozous koopmani* specimens in Orihuela et al., forthcoming). By then Amerindians, both pre-Arawak “archaic” and Arawaks (Taino) were already well-established in the area (Tabío and Rey, 1979; Roksandic et al., 2015; Chinique et al., 2016), thus human-caused forest fires cannot be ruled out. Although, based on the lack of archaeological evidence, we consider this as a result of a natural fire on the upper escarpment of the hill. Natural fires are commonly ignited by lightning, as is the case in Cuba (Medina and Alfonso, 2000; Ramos, 2002). Microcharcoal deposits in Cuba (Jiménez et al., 2005) and other parts of the Greater Antilles, although common in some cave and lacustrine deposits, have been difficult to attribute to human action (Burney et al., 1994; Haug et al., 2001; Lane et al., 2013; Caffrey and Horn, 2014). Furthermore, we did not find any archaeological evidence (e.g., tool cut marks) at the doline deposit that could suggest human involvement in these localized fires.

## CONCLUSIONS

The deposit reported and interpreted here from Cueva de los Nesofontes, in northeastern Mayabeque province, provides a rich source of biogeographical and paleoecological information with which to understand the pre-Columbian (Amerindian) environmental history of the late Holocene of Cuba. Through a multidisciplinary and multiproxy approach, we access the formational history and source of the deposit, including the survivorship and coexistence of fauna on a millennial-scale. From these data, we infer that the cone deposit in the main doline gallery is a secondary repository of primarily amalgamated, multisource deposit located in the upper levels of the main sinkhole. Although the deposit is mostly pellet-derived, it was slightly mixed over time with organisms by sedimentation and reworking.

The stratigraphic architecture (disconformities and erosional surfaces) that suggest flooding events helped transport sediments and organic remains into the deposition cone below the sinkhole. The deposition was controlled by the slope’s incline and is marked by a slow sedimentary regime (slow sedimentary rates), suggesting that it was slow to form and time-averaged. The oxygen isotopes suggest a change in the tendency from wet and warm conditions during BC 40 – AD 90 to slightly wetter and warmer conditions thereafter. A positive excursion was registered during AD 660–770, before the medieval warm period, that suggests drier and colder local conditions, which does not agree with most other circum-Caribbean records.

The radiocarbon dates yielded by faunal bone collagen indicate that the sampled portion of the deposit is less than 2000 years old, and thus within the pre-Columbian Amerindian interval of the Late Holocene. Direct radiocarbon dates from extinct fauna provide last occurrence dates for the extinct fruit bats *Artibeus anthonyi* and *Phyllops vetus*, plus the extinct island-shrew *Nesophontes major*, previously without direct LAD dates. These dates support the inference that some Cuban extinct land mammal taxa, formerly believed to have disappeared during the late Pleistocene-early Holocene, survived well into the late Holocene, and several thousands of years of human presence in the archipelago (MacPhee et al., 1999; Jiménez et al., 2005; Orihuela, 2010; Orihuela and Tejedor, 2012). The association of these species within the dated intervals of the deposit also provides new records supporting a wider distribution for species that are either extinct or severely endangered, such as the bats *Natalus primus* and *Antrozous koopmani*. Other taxa, such as *Psittacara eups*, *Corvus*, *Colaptes,* and *Solenodon cubanus* are locally extinct or no longer occur in the region, but their presence in the deposit support their existence in the surrounding habitats up to the very late Holocene.

The integration of isotopic data as a proxy for dietary preferences with evidence of niche partitioning may serve to better elucidate unexplored causes of extinction of Antillean land mammals, which we have only recently begun to explore (e.g., Cooke and Crowley, 2018; this work). The information that can be gleaned with simultaneous analyses of assemblage structure and resource partitioning can help elucidate aspects of competition and trophic guilding, which when coupled to climatic and anthropogenic factors, can provide a more naturalistic (realistic) explanation to the asynchronous and taxon-specific extinction of land vertebrates during the Antillean Late Holocene.

